# Chromosome-level genome assemblies and annotations of Amaranthus spinosus, Amaranthus acanthochiton, Amaranthus arenicola, and Amaranthus floridanus

**DOI:** 10.64898/2026.08.21.746229

**Authors:** Damilola A. Raiyemo, Isabel Werle Noe, Ramandeep Kaur, Lauren Whitt, Sarah B. Carey, Haley Hale, Keyana J. Lewis, Lauren Womack, Alex Harkess, Victor Llaca, Kevin Fengler, Eric L. Patterson, Todd A. Gaines, Patrick J. Tranel

**Affiliations:** Department of Crop Sciences, University of Illinois, Urbana, Illinois, USA; HudsonAlpha Institute for Biotechnology, Huntsville, Alabama, USA; Genome Center of Excellence, Corteva Agriscience, Johnston, Iowa, USA; Department of Plant, Soil and Microbial Sciences, Michigan State University, East Lansing, Michigan, USA; Department of Agricultural Biology, Colorado State University, Fort Collins, Colorado, USA

**Keywords:** Amaranthaceae, comparative genomics, dioecy, sex chromosome, weed

## Abstract

*Amaranthus* L. spans aggressive agricultural weeds, ornamentals, and ancient pseudocereals. Species within the genus vary in morphology, environmental tolerance, and sexual systems, making them well-suited for studying reproductive evolution and plant adaptation. To investigate sex chromosome architecture within the genus, we generated chromosome-level assemblies of a monoecious amaranth (*Amaranthus spinosus*) and three dioecious species (*A. acanthochiton, A. arenicola*, and *A. floridanus*) using PacBio high-fidelity (HiFi) long reads. We paired these data with Dovetail Genomics Omni-C sequencing to achieve haplotype resolution for *A. spinosus* and *A. acanthochiton*, and we used reference-guided scaffolding for the remaining two species. The assemblies are highly contiguous, with sizes ranging from 394.24–607.10 Mbp, contig N50 from 0.63–8.76 Mbp, and scaffold N50 from 22.44–37.97 Mbp. Evaluation of the assemblies and annotations revealed 96.3–97.6%, and 97.6–98.3% BUSCO completeness, respectively. Comparative genomic analysis revealed that the Chromosome 1 inversions and Robertsonian fusion previously reported in *A. tuberculatus* are conserved in *A. acanthochiton* and consistent with the architecture of *A. arenicola* and *A. floridanus*, suggesting that the evolution of dioecy in this clade predates subsequent speciation. In parallel, multiple homologs of *Rf1* on Chromosome 3 of *A. spinosus*, a monoecious species that exhibits spatial separation of male and female flowers and is closely related to the dioecious *A. palmeri*, were identified. Together, this study provides foundational resources for advancing evolutionary, ecological, and agronomic research across the genus, including herbicide resistance evolution and weediness traits.

**PLAIN LANGUAGE SUMMARY:** The genus *Amaranthus* comprises agronomically important species, including troublesome weeds, ornamentals, grain crops, and leafy vegetables, which exhibit different sexual systems. We present high-quality, chromosome-level genomes and annotations for four *Amaranthus* species: *A. spinosus*, a widespread agricultural weed, alongside three wild relatives (*A. acanthochiton, A. arenicola*, and *A. floridanus*). These assemblies provide valuable resources for accelerating research into how species within the genus adapt to different environments and evolve traits such as herbicide resistance.

## 1 INTRODUCTION

The genus *Amaranthus* L. comprises approximately 70 – 80 species and has its primary diversity in the Americas, with a cosmopolitan distribution (Sauer, 1967; Trucco et al., 2005). Species within the genus are of notable ecological, cultural, and agricultural importance, ranging from weedy invaders to ornamentals and ancient pseudocereals domesticated by pre-Columbian societies in Mesoamerica (Adhikary & Pratt, 2015; Stetter & Schmid, 2017). This ecological breadth, together with morphological and reproductive variation, positions the genus as a compelling model for studies of sexual-system evolution, reproductive development, and plant adaptation.

*Amaranthus* species exhibit two major sexual systems: monoecy (separate male and female flowers on the same individual) and dioecy (male and female flowers on separate individuals) (Eliasson, 1988; Sauer, 1955). Monoecy is most common among the cultivated grain species and several weeds within the genus (Bayón, 2022; Mallory et al., 2008). An exception within commonly discussed monoecious species is *Amaranthus spinosus*, which typically bears predominantly staminate flowers in terminal spikes and pistillate flowers in axillary clusters, reflecting spatial separation of reproductive organs within individuals (Waselkov et al., 2018). *A. spinosus* is a widely distributed weed species throughout tropical and subtropical regions (Bayón, 2022; Sarker & Oba, 2019). Characterized by spiny structures in the leaf axils and a propensity for disturbed habitats, it has been well studied in weed science (Chauhan & Abugho, 2012; Chauhan & Johnson, 2009; Nandula et al., 2014; Odero & Wright, 2022), though its genetic and reproductive biology have been investigated far less intensively.

Conversely, dioecy characterizes a subset of species, including the agriculturally important *A. tuberculatus* and *A. palmeri*, and ecologically important taxa such as *A. acanthochiton, A. arenicola* and *A. floridanus* (Iamonico, 2020; Raiyemo & Tranel, 2023; Sauer, 1972; Waselkov et al., 2018). *Amaranthus acanthochiton*, commonly known as greenstripe amaranth, is native to arid and semi-arid regions of the southwestern United States and northern Mexico, and was traditionally used by the Hopi as food (Minnis, 1991; Sauer, 1955, 1957). Despite its cultural importance and xeric adaptation, it remains comparatively understudied. *Amaranthus arenicola* is native to the central and southwestern Great Plains (J. D. Sauer, 1955, 1972), and has been recorded as introduced in Hawai’i, Korea, and China, occurring on sandy substrates, along river and lake margins, and in disturbed sites, including agricultural fields (Faccenda & Ross, 2024; Xu et al., 2022). *Amaranthus floridanus* is a Florida endemic of very limited range, restricted to wet coastal areas of the peninsula, with morphological and genomic evidence supporting a very close relationship to *A. tuberculatus* (Raiyemo et al., 2023; Raiyemo & Tranel, 2023; Sauer, 1957; Waselkov et al., 2018). The co-occurrence of monoecy and dioecy within the genus provides a framework for dissecting the genetic and evolutionary drivers of transitions between sexual systems (Timerman et al., 2026).

Cytogenetic and, more recently, genomic studies have contributed to our understanding of the evolution and structure of sex-determining regions (SDRs) in dioecious *Amaranthus* species (Grant, 1959; Kreiner et al., 2025; Montgomery et al., 2021; Raiyemo, Montgomery, et al., 2025). Evidence of male heterogamety (XY) has been established in *A. tuberculatus* and *A. palmeri*, as well as identification of male-specific Y contigs (Montgomery et al., 2019, 2021). Furthermore, recent evaluations of the relationships among dioecious species and the genomic landscape of SDRs have revealed distinct architectures between these two weed species (Raiyemo, Cutti, et al., 2025; Raiyemo et al., 2023; Raiyemo, Montgomery, et al., 2025). In *A. tuberculatus*, chromosome-level comparative analyses localize the candidate SDR to the middle of Chromosome 1 (Kreiner et al., 2025; Raiyemo, Cutti, et al., 2025), while in *A. palmeri* the candidate SDR resides at the distal end of Chromosome 3 (Raiyemo, Montgomery, et al., 2025). The non-syntenic nature of these SDRs, together with phylogenetic evidence, supports at least two independent origins of dioecy within the genus, corresponding to two distinct clades: (i) one in which the dioecious *A. palmeri* is closely related to the monoecious *A. spinosus*; and (ii) the other including *A. tuberculatus* together with *A. acanthochiton, A. arenicola*, and *A. floridanus* (Raiyemo, Cutti, et al., 2025; Raiyemo et al., 2023; Raiyemo, Montgomery, et al., 2025; Timerman et al., 2026). The SDR architecture characterized in *A. tuberculatus*, however, has not yet been assessed across the other dioecious species within the genus. More broadly, the determinant sex factors and the molecular underpinnings of floral initiation and development across sexual systems remain incompletely resolved. Comparative analyses spanning multiple species are essential to identify conserved genes and characterize genus-wide patterns of SDR differentiation.

Here, using chromosome-level assemblies and comparative genomics of *A. spinosus, A. acanthochiton, A. arenicola*, and *A. floridanus*, we elucidate conserved and divergent features of sex chromosomes across the genus. Building on prior foundational work, our study refines and expands current understanding of reproductive evolution in *Amaranthus*.

## 2 MATERIALS AND METHODS

### 2.1 Plant materials, library preparation, and sequencing

Species were obtained from the USDA Germplasm Resources Information Network (GRIN) (Table S1). Seeds from each were grown in pots filled with a growing media that included Sunshine LC1 (Sun Gro Horticulture, 770 Silver Street Agawam, MA) growing mix, soil, peat, and torpedo sand (3:1:1:1 by weight). Young leaves were harvested from each species following visible flower formation on the spike of inflorescence. Harvested leaves were flash frozen in liquid nitrogen and stored in −80 C pending DNA extraction. High molecular weight (HMW) DNA was extracted from one individual of *A. spinosus* and one male individual of *A. acanthochiton* using the NucleoBond HMW DNA extraction kit (Macherey-Nagel, Düren, Germany) with 1 g of young leaf material as input, following the manufacturer’s recommendations, and sized on a Femto Pulse (Agilent Technologies, Santa Clara, CA). The isolated HMW DNA samples were used for library preparation, and the libraries were sequenced on a PacBio Revio. Shotgun genomic libraries were also prepared for *A. acanthochiton* using the seqWell purePlex DNA library preparation kit (SeqWell, Beverly, MA), and the libraries sequenced paired-end (2 x 150) on Illumina NovaSeq X plus. Sequencing was carried out at HudsonAlpha Institute for Biotechnology (Huntsville, Alabama). The same tissues for which

HMW DNA were extracted for PacBio HiFi sequencing were also used for Omni-C library preparation and sequencing (Dovetail Genomics, Scotts Valley, California) using 1 g of input material. Collected leaf tissues from one male and one female of each species of *A. arenicola* and *A. floridanus* were also flash-frozen in liquid nitrogen and shipped to the Genome Center of Excellence at Corteva Agriscience for HMW DNA extraction, library preparation and sequencing. PacBio HiFi reads ranging from 10.74 – 84.22 Gb (average read length 9.56 – 20.25 kb) with estimated coverage ranging from 18 – 135X were used for genome assembly (Table S2).

### 2.2 Genome assembly, repetitive element analysis, and annotation

#### 2.2.1 Assembly and scaffolding

PacBio HiFi reads for *A. acanthochiton* and *A. spinosus* were screened for adapter contamination with HiFiAdapterFilt v3.0.0 (Sim et al., 2022) and quality checked with Nanoplot v1.44.1 (De Coster & Rademakers, 2023). The nuclear genomes were assembled with HiFiasm v0.25.0 (Cheng et al., 2021), integrating Omni-C paired-end data for phasing. Due to the high homozygosity of the *A. spinosus* genome, HiFiasm was run with the ‘--hom-cov’ parameter set to 147. Both the *A. acanthochiton* and *A. spinosus* assemblies were polished with Racon v1.5.0 (Vaser et al., 2017) to correct residual indel errors observed in homopolymeric regions of HiFiasm contigs, and short contigs less than 30 kb (*A. acanthochiton*) and 40 kb (*A. spinosus*) were removed with BBmap v39.19 (Bushnell, 2014). The polished assemblies were then screened for contaminant sequences using NCBI’s FCS-GX tool v0.5.5 (Astashyn et al., 2024). Omni-C paired-end reads for each species were mapped to the putative haplotypes from their respective HiFiasm assembly with BWA-MEM v0.7.19 (Li, 2013) using default settings, with the additional parameter ‘-5SP’. The Omni-C alignments were filtered to remove duplicate read-pairs using pairtools v1.1.3 (Open2C et al., 2024). The filtered alignments were then scaffolded with YaHS v1.2.2 (Zhou et al., 2023) and contact maps were generated with YAHS’s built-in juicer tools (Dudchenko et al., 2018). Juicebox v1.11.08 (Durand et al., 2016) was used for visualization and to manually edit the contact maps. Paired-end reads for male and female *A. acanthochiton* individuals were used to confirm haplotype phasing of the sex chromosomes following the methods in Carey et al. (2024). Briefly, Meryl v1.4.1 (Rhie et al., 2020) was used to count 21-mers in each individual and return *k*-mers shared across all male individuals that were not found in any female individuals. The male-specific *k*-mers were then mapped to the *A. acanthochiton* assembly using BWA-MEM (Li, 2013) with parameters ‘-k 21 -T 21 -a -c 10’. The mapping coverage was calculated with SAMtools v1.19.2 (Danecek et al., 2021) and confirmed all Y-linked contigs were phased into Hap1 and scaffolded together.

The raw data for *A. arenicola* and *A. floridanus* males were also assembled using Hifiasm v0.25.0 (Cheng et al., 2021) with parameters: --primary, --telo-m TTTAGGG --telo-d 60000. Due to the unavailability of Omni-C data for both species, the primary and alternate contig assemblies were combined within each species and scaffolded to chromosome levels using RagTag v2.1.0 (Alonge et al., 2022), guided by a closely related, haplotype-resolved *A. tuberculatus* genome (Raiyemo, Cutti, et al., 2025) to facilitate phasing. All parameters in RagTag were default, except that minimap2 -x asm20 parameter was used during the correction and scaffolding steps to account for divergence between *A. tuberculatus* and the two species. Similarly, the raw HiFi reads for the females of both *A. floridanus* and *A. arenicola* were assembled as described above, and the combined primary and alternate contig assemblies scaffolded using the Hap2 of *A. tuberculatus* genome. Combining primary and alternate contigs yielded 1,054.9–1,135.1 Mbp per assembly of which 53.2–55.6% (585.1–606.8 Mbp) was placed during scaffolding and retained. Unplaced contigs after scaffolding for the four genome assemblies were discarded and not considered for downstream analyses. These discarded unplaced sequences were mainly short, with mean lengths of 30.2 – 45.3 kb. Plastomes and mitogenomes for each of the species were assembled and annotated with OatK v1.0 (Zhou et al., 2025) using the raw HiFi reads described above.

#### 2.2.2 Repetitive element analysis

Species-specific repeats were first determined for each of the four assembled genomes using RepeatModeler v2.0.7 (Flynn et al., 2020), including its integrated LTR structural annotation module (-LTRStruct). The species-specific libraries were combined each with RepBase database (RepeatMaskerEdition-20181026), and a RepeatMasker-derived “viridiplantae” repeats library that had been clustered with CD-HIT-EST v4.8.1 to reduce redundancy. The nonredundant libraries were then used to annotate repeats in each of the assemblies using RepeatMasker v4.2.2 with parameters -s -gff -lib -e rmblast -xsmall -a (http://www.repeatmasker.org/RepeatMasker/). The annotated repeats for each of the assemblies were summarized using the Perl script “buildSummary.pl” within RepeatMasker. To refine LTR annotations and determine LTR assembly index (LAI), each genome assembly was further analyzed using LTR_retriever v3.0.7 (Ou & Jiang, 2018). Telomere repeat motif (TTTAGGG)_n_, where n = 4, was used to determine if telomeres were contiguously assembled in the genomes. The motif was searched in each assembly using BLASTN (Camacho et al., 2009) with -task blastn-short parameter.

#### 2.2.3 Gene prediction and functional annotation

Genes in the assembled genomes of *A. spinosus, A. acanthochiton, A. arenicola*, and *A. floridanus* were predicted using BRAKER3 long reads version (Gabriel et al., 2024; Huang & Li, 2023; Li, 2023). Clustered Iso-Seq reads of *A. tuberculatus* from previously published dataset (Raiyemo, Cutti, et al., 2025) were mapped separately to all haplotype assemblies using minimap2 with parameters: -ax splice:hq -uf --secondary=no. The aligned transcripts and the clade-partitioned OrthoDB v12 (Viridiplantae) (Tegenfeldt et al., 2025), combined with protein sequences of *A. hypochondriacus* v2 and *A. cruentus* were used as evidence for BRAKER3. Primary isoforms (i.e., longest CDS) were selected from the gene models using the AGAT tool, *agat_sp_keep_longest_isoform.pl*. PASA was then utilized to update structural gene annotations, particularly by adding 5’ and 3’ UTRs and refining exon-intron boundaries. Full-length transcripts were first aligned to the respective haplotype assemblies with minimap2 using the *Launch_PASA_pipeline.pl* script with the following options in the alignAssembly.txt configuration file: validate_alignments_in_db.dbi:--MIN_PERCENT_ALIGNED=90; validate_alignments_in_db.dbi:--MIN_AVG_PER_ID=95; validate_alignments_in_db.dbi:--MAX_INTRON_LENGTH 10000; subcluster_builder.dbi:-m=50. The initial annotation set from BRAKER after selecting longest isoform with AGAT were supplied to PASA for comparison and model updating. Functional annotation was performed using EnTAP v2.3.0 (Hart et al., 2020), and transfer RNA (tRNA) genes were predicted with tRNAscan-SE-2.0.12 (Chan et al., 2021). Postprocessing steps, including merging the PASA output, EnTAP annotations, and tRNA predictions, as well as generating a final gff3 annotation file were performed using custom scripts (https://github.com/Alexdami17/Amaranthus-genome-assembly). Assembly completeness was evaluated using BUSCO v6.0.0 (Simão et al., 2015) in genome mode, and annotation completeness was evaluated in protein mode on the primary-isoform proteomes with embryophyta_odb10 database. Genome characteristics were determined using “*agat_sp_statistics.pl*” from AGAT Toolkit v1.4.1 (Dainat, 2022).

### 2.3 Comparative analyses and sex chromosome architecture

Synteny analyses among the haplotype assemblies (*A. spinosus, A. acanthochiton, A. arenicola*, and *A. floridanus*) from this study and available chromosome-level assemblies of *Amaranthus* species: *A. hypochondriacus* (Lightfoot et al., 2017), *A. cruentus* (Ma et al., 2021), *A. tricolor* (Wang et al., 2023), *A. palmeri, A. retroflexus, A. hybridus* (Raiyemo, Montgomery, et al., 2025), and *A. tuberculatus* (Raiyemo, Cutti, et al., 2025) were performed using GENESPACE v1.2.3 with default parameters (Lovell et al., 2022). Riparian plots were generated using the “*plot_riparian*” function in GENESPACE. Chromosomes were ordered to maximize synteny with the *A. tricolor* as reference, and ribbons were color-coded to indicate the sex chromosomes. Haplotype assemblies of each species were further aligned using Minimap2 v2.28-r1209 (Li, 2018) and the alignments visualized using D-Genies (Cabanettes & Klopp, 2018).

The alignments were further processed with SyRI (Goel et al., 2019) to identify structural rearrangements on putative sex chromosomes, which were then plotted using plotsr (Goel & Schneeberger, 2022). To identify genes conserved across species within the region associated with sex in *A. tuberculatus* (Chr01: 14.02 – 45.81 Mbp), syntenic gene counterparts were extracted for *A. acanthochiton, A. arenicola*, and *A. floridanus* using GENESPACE, and genes with non-missing assignments in GENESPACE were considered syntenically conserved within the interval. Conserved genes that may have been displaced outside the focal region due to rearrangements, assembly, or annotation differences were further investigated using protein-level searches in MMseqs2 (easy-search) with the following parameters: --min-seq-id 0.50 -c 0.60 --cov-mode 1 -e 1e-10 --max-seqs 1. Results from GENESPACE synteny and MMseqs were integrated to generate a presence-absence matrix. Genes supported by either pipeline were retained, while those lacking evidence across all target species were excluded.

To find the 37 genes, including *Rf1* and *TLC* previously reported, within the MSY region of *A. palmeri* in *A. spinosus*, protein sequences from *A. palmeri* were searched against the protein sequences from *A. spinosus* using MMseqs easy-search with parameters: --min-seq-id 0.30 -c 0.60 --cov-mode 1 -e 1e-5 --max-seqs 30. The protein sequences were then aligned using MAFFT v7.511 (Katoh & Standley, 2013), and the aligned sequences trimmed using trimAL (Capella-Gutiérrez et al., 2009) with the -gappyout parameter. Phylogenetic tree was constructed with the trimmed sequences using IQ-TREE v3.0.1 (Wong et al., 2026) with parameters: -m MFP -bb 1000 -alrt 1000 -nt AUTO. The resulting consensus tree was visualized and annotated with FigTree v1.4.5_pre (https://tree.bio.ed.ac.uk/software/figtree/).

### 2.4 Change-point analysis of synonymous divergence (K_S_)

Protein sequences from the X-and Y-haplotype assemblies of *A. acanthochiton*, along with previously reported assemblies of *A. palmeri* and *A. tuberculatus*, were compared using reciprocal best-hit (RBH) searches implemented in MMseqs2 (Steinegger & Söding, 2017). To minimize the inclusion of paralogous matches arising from gene duplication or translocation events, RBH pairs were further filtered to retain only strict one-to-one pairs and gene pairs located on homologous chromosomes (i.e., both gene identifiers corresponding to the same chromosome number in their respective haplotype assemblies). These filtered one-to-one homologous X-Y gene pairs were treated as putative gametologs and aligned using MAFFT v7.511 (Katoh & Standley, 2013). Codon-based nucleotide alignments were generated using PAL2NAL v14 (Suyama et al., 2006), and synonymous divergence (Ks) values were estimated from the nucleotide alignments using Ka/Ks Calculator 3.0 (Zhang, 2022). Generalized Additive Models (GAM) with a robust scaled-t error distribution were first fitted to the Ks values using penalized regression splines (k = 30 basis functions) implemented in the R package *mgcv*.

Bayesian changepoint models were then fit separately using the R package *mcp* to infer potential evolutionary strata. Prior to modeling, extreme Ks values were identified using Rosner’s generalized ESD test for multiple outliers, as implemented in the R package *EnvStats*, and detected outliers were removed. Throughout the analysis, gametolog positions were ordered according to their coordinates on the X chromosome (Hap2 assembly) to provide a consistent spatial framework for changepoint inference. Bayesian models assuming piecewise constant Ks means were fit with zero to three changepoints (corresponding to one to four plateaus). Each model was run for 100,000 MCMC iterations across four chains, following a burn-in (adaptation) period of 20,000 iterations. MCMC convergence was assessed via the Gelman-Rubin R-hat statistic (threshold R-hat < 1.1) and effective sample size. Competing models were compared using leave-one-out cross-validation (LOO), and the pseudo-Bayesian model averaging (pseudo-BMA) weights derived from LOO cross-validation, both implemented in the *loo* package (Vehtari et al., 2017). Where models allowing multiple changepoints failed to converge, simplified one-changepoint models were fitted as well.

### 2.5 Gene-centric pangenome analysis

Protein sequences of 11 *Amaranthus* species, including *A. hypochondriacus* (Lightfoot et al., 2017), *A. cruentus* (Ma et al., 2021), *A. tricolor* (Wang et al., 2023), *A. palmeri* (Raiyemo, Montgomery, et al., 2025), *A. hybridus* (Raiyemo, Montgomery, et al., 2025), *A. retroflexus* (Raiyemo, Montgomery, et al., 2025), *A. tuberculatus* (Raiyemo, Cutti, et al., 2025), *A. spinosus, A. acanthochiton, A. arenicola*, and *A. floridanus*, representing 17 genome assemblies (including 12 haplotype assemblies), were used to construct a gene-centric pangenome. Orthology relationships were inferred using OrthoFinder v2.5.4 with default parameters (Emms & Kelly, 2019). The orthogroup gene counts per genome (Orthogroups.GeneCount.tsv) were used to generate a presence–absence matrix, with an orthogroup considered present when ≥1 gene was assigned to that orthogroup. To assess the effect of haplotype-resolved assemblies, pangenome analyses were conducted both at the assembly level and after collapsing Hap1 and Hap2 to the species level, with an orthogroup considered present in a species when detected in either haplotype. Orthogroup occupancy was calculated as the number of taxa in which an orthogroup was present divided by the total number of taxa included in the analysis. Following the general occupancy-based partitioning of pangenome components (Kaur et al., 2024; Tettelin & Medini, 2020), integer occupancy thresholds were used to accommodate the discrete distribution of orthogroup frequencies across the 11 taxa examined. Orthogroups were classified as core when present in all 11 taxa, soft-core when present in 10 taxa, shell when present in 3–9 taxa, cloud when present in two taxa, and private when present in a single taxon. Pangenome accumulation curves were generated separately for the assembly-level and species-level analyses by randomly permuting the order of genomes or collapsed species, respectively, 500 times and iteratively adding one at each step. At each step, the cumulative number of observed orthogroups (pan) and the number of orthogroups shared by all genomes or species included (core) were recorded. Curves were summarized as the mean with percentile bands across permutations.

## 3 RESULTS

### 3.1 Genome assembly and annotation metrics, and repeat landscape

We assembled the genomes of four *Amaranthus* species, whose representative morphologies are shown in Figure 1: two haplotypes for *A. spinosus* (17 chromosomes per haplotype), a male *A. acanthochiton* (16 chromosomes), and male and female individuals of *A. arenicola* and *A. floridanus*. The latter two were each assembled into contigs and scaffolded into chromosome levels using *A. tuberculatus* Hap1 (containing the Y chromosome) or Hap2 (containing the X chromosome) assembly (Table S1).

**FIGURE 1.**
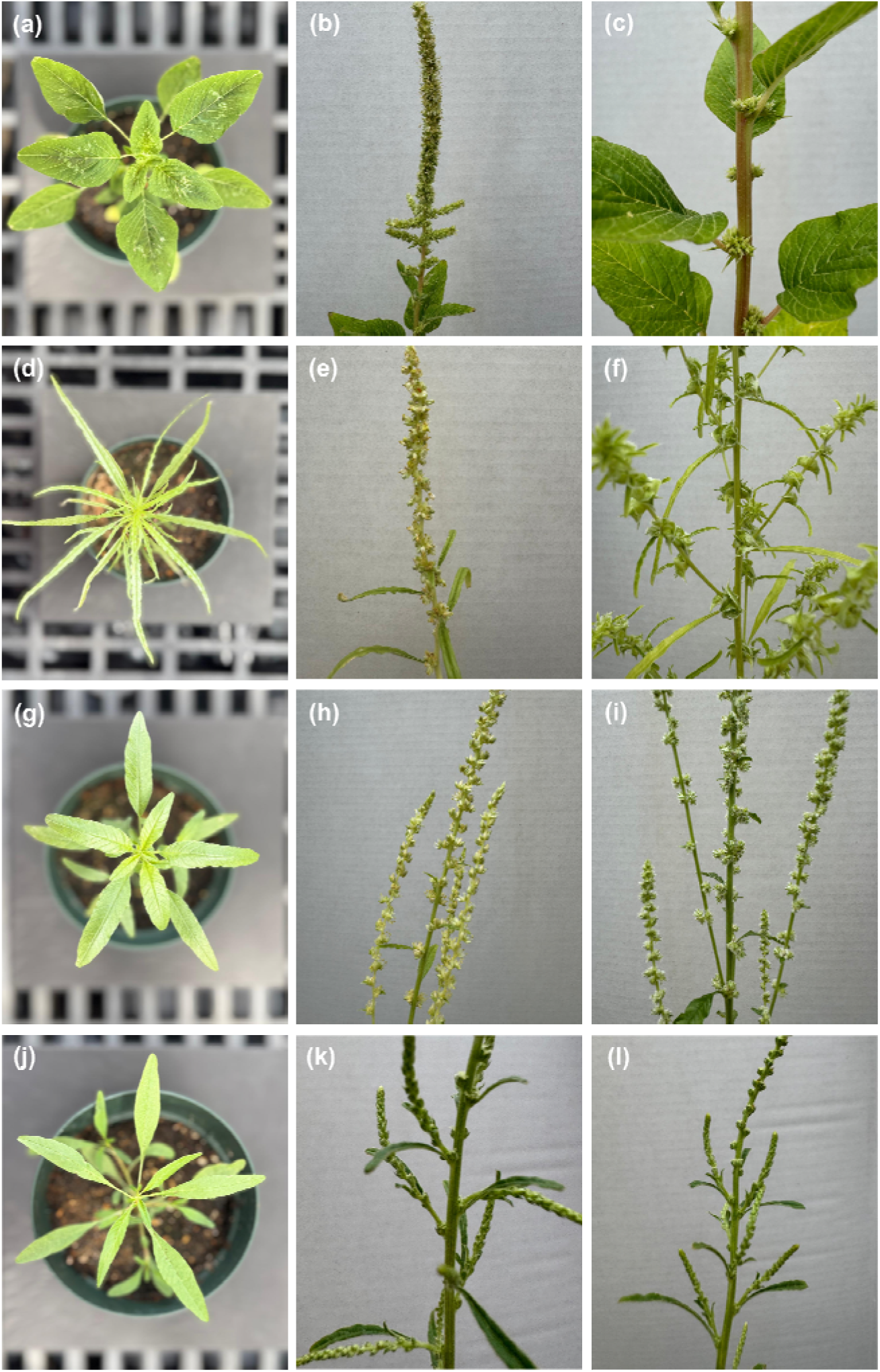
Representative vegetative and reproductive morphology across four *Amaranthus* species. Monoecious *Amaranthus spinosus* (a – c): (a) vegetative growth; (b) male inflorescence; (c) female reproductive structures in axils. Dioecious *A. acanthochiton*: (d – f), *A. arenicola* (g – i), and *A. floridanus* (j – l) vegetative growth (left) and male (center) and female (right) inflorescences.

Hence, throughout the manuscript, *A. tuberculatus*-Hap1-scaffolded assemblies for *A. arenicola* and *A. floridanus* are referred to as Hap1 while *A. tuberculatus*-Hap2-scaffolded assemblies are referred to as Hap2 (Table S1). For *A. acanthochiton*, a male individual was assembled and scaffolded *de novo*; chromosomes were however named to match the corresponding homologs in *A. tuberculatus*. Assembled genome sizes were compared with previously reported estimates for each species: flow cytometry estimate for *A. spinosus* (Stetter & Schmid, 2017) and our earlier *k*-mer-based estimates (Raiyemo et al., 2023) for the remaining three species (Table 1 and Table S2).

**TABLE 1.**
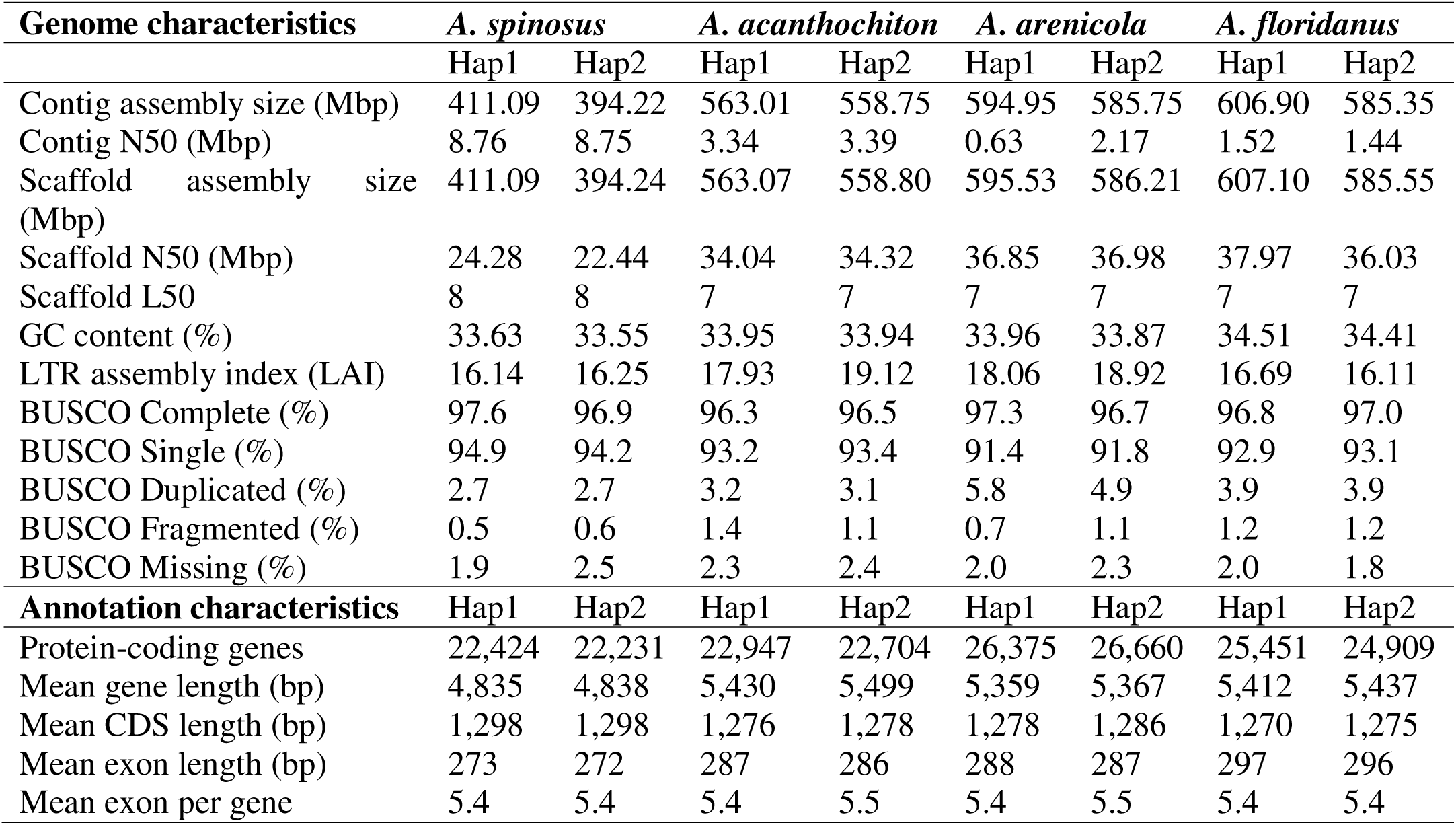
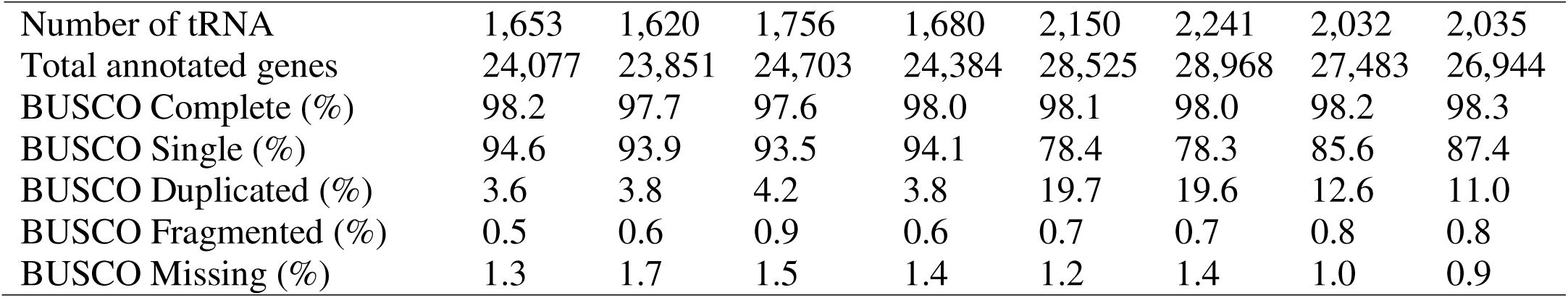
Comparison of nuclear genome assembly statistics among *Amaranthus spinosus, Amaranthus acanthochiton, Amaranthus arenicola*, and *Amaranthus floridanus*.

The *A. acanthochiton* assemblies represent 89.9–90.6% of the corresponding estimate, whereas those of *A. arenicola* and *A. floridanus* approach or exceed the *k*-mer-based estimates (98.2 – 105.8%), consistent with retained allelic redundancy. Genome assembly completeness ranged from 96.3 to 97.6% complete BUSCOs across the species based on embryophyta_odb10 database. Protein annotation completeness was similarly high, revealing 97.6 to 98.3% of the reference orthologs (Table 1). This marginal increase in annotation completeness arises from differences in how gene prediction is handled: genome mode relies on BUSCO’s internal *ab initio* prediction, whereas protein mode is applied to evidence-based gene models.

Analysis of repetitive elements revealed 62.61–69.73% of the assemblies were made up of repeats (Table S3). The LTR retrotransposon (specifically, *Ty3* and *Copia* elements) constituted a high proportion of repeats in the genome assemblies (Table S3). RepeatMasker repeat landscapes also revealed differences in the age structure of transposable elements (TE) across the genomes, particularly when comparing *A. spinosus* and *A. palmeri* with the more distantly related *A. acanthochiton, A. arenicola, A. floridanus*, and *A. tuberculatus* (Figure 2).

**FIGURE 2.**
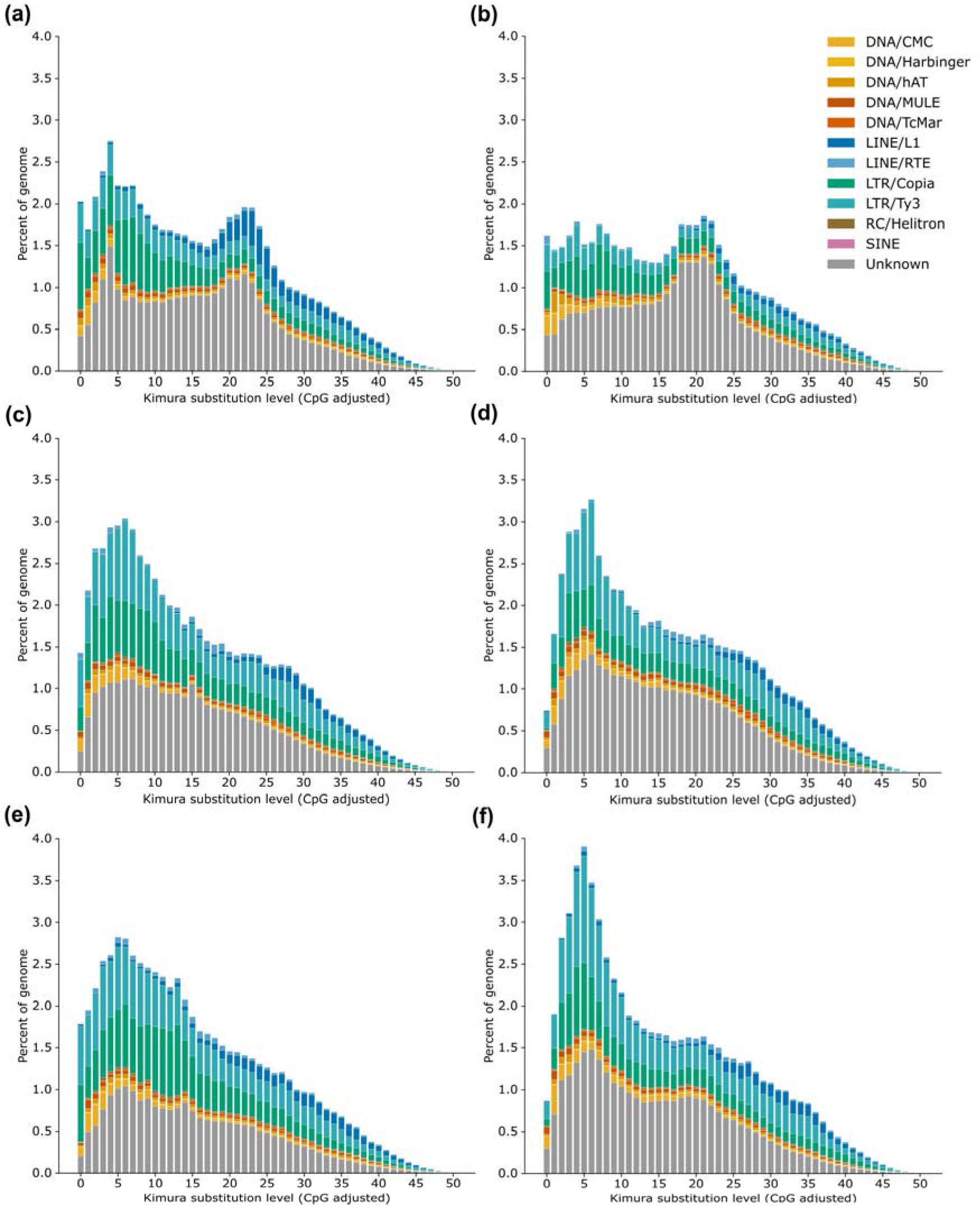
Repeat landscapes for a) *A. spinosus*, b) *A. palmeri*, c) *A. acanthochiton*, d) *A. arenicola*, e) *A. floridanus*, and f) *A. tuberculatus* from RepeatMasker. The y-axis shows the percentage of the genome occupied, and the x-axis shows the kimura-2 substitution level (CpG adjusted). Color codes indicate the different repeat superfamilies.

The distribution of Kimura 2-parameter divergence values showed TEs were concentrated in intermediate bins (18 – 24) for *A. spinosus* and *A. palmeri*, consistent with older insertion events. Distinct peaks within these intervals correspond to lineage-specific expansions of LINE and LTR elements. In contrast, *A. acanthochiton, A. arenicola, A. floridanus*, and *A. tuberculatus*, exhibited peaks at k < 10, indicating recent transposition activity. These low-divergence peaks were primarily associated with LTR elements (Figure 2). Telomere analysis revealed the presence of telomeric repeats at the ends of most chromosomes (ranging from 59 to 91% of possible telomeric ends) for both Hap1 and Hap2 assemblies of each species (Tables S4A – S4H).

### 3.2 Comparative analyses and sex chromosome architecture

Self-alignment of the genomes reveals single, continuous diagonals without breaks, gaps, or off-diagonal matches, indicating no large-scale structural misassembly (Figures S1 – S10). At finer scales, however, the *A. arenicola* and *A. floridanus* assemblies retain allelic redundancy introduced by combining primary and alternate contigs prior to scaffolding: 11.0–19.7% of BUSCOs are duplicated in their annotations, compared with 3.6–4.2% in *A. spinosus* and *A. acanthochiton*. These duplicate copies are typically separated by a few hundred kilobases, consistent with adjacent placement of allelic contigs. Alignment between *A. spinosus* and the closely related *A. palmeri* shows broad collinearity between the assemblies, with four visible localized inversions on Chromosomes 1, 2, 13, and 17, suggesting subtle structural differences between the genomes (Figure S2). Chromosome 3 appears to be largely collinear between the two species, with notable “micro-inversions” (Figure S11). Alignment of *A. arenicola* and *A. floridanus* Hap1 or Hap2 to the corresponding *A. tuberculatus* Hap1 or Hap2 assemblies to which they were scaffolded also revealed high synteny between the assemblies (Figures S5, S6, S8, and S9). Assessment of inversions revealed 341 inversions (Hap1 vs. Hap2) across the 16 chromosomes of *A. acanthochiton*, 166 in *A. arenicola*, and 141 in *A. floridanus* (Table S5 – S7). Across the three species, the two largest inversions were consistently located on Chromosome 1. Notably, because *A. arenicola* and *A. floridanus* assemblies were ordered and oriented against different *A. tuberculatus* haplotypes, inversions spanning a scaffolding junction are not independent of the reference. Therefore, restricting inversions to those contained within a single contig in both haplotypes retains 132 of 166 inversions in *A. arenicola* and 106 of 141 in *A. floridanus*. Also, the proportion of inversions retained falls steeply with size. In *A. arenicola*, all 95 inversions below 10 kb and 28 of 33 between 10 and 100 kb are within contigs; however, 21 of 29 between 100 kb and 1 Mb and 8 of 9 above 1 Mbp span scaffolding junctions. Similarly, for *A. floridanus*, all 58 inversions below 10 kb and 33 of 38 between 10 and 100 kb are within contigs, while 16 of 31 between 100 kb and 1 Mbp and all 14 above 1 Mbp span scaffolding junctions. The two largest inversions on Chromosome 1 for both species fall into the >1 Mbp category. Inversion counts for both *A. arenicola* and *A. floridanus* should therefore be regarded as lower bounds and not directly comparable with that reported for *A. acanthochiton*.

Synteny analysis revealed that the fusion of two ancestral chromosomes into a single unit, previously reported for Chromosome 1 of *A. tuberculatus*, is conserved across the three species (Figure 3).

**FIGURE 3.**
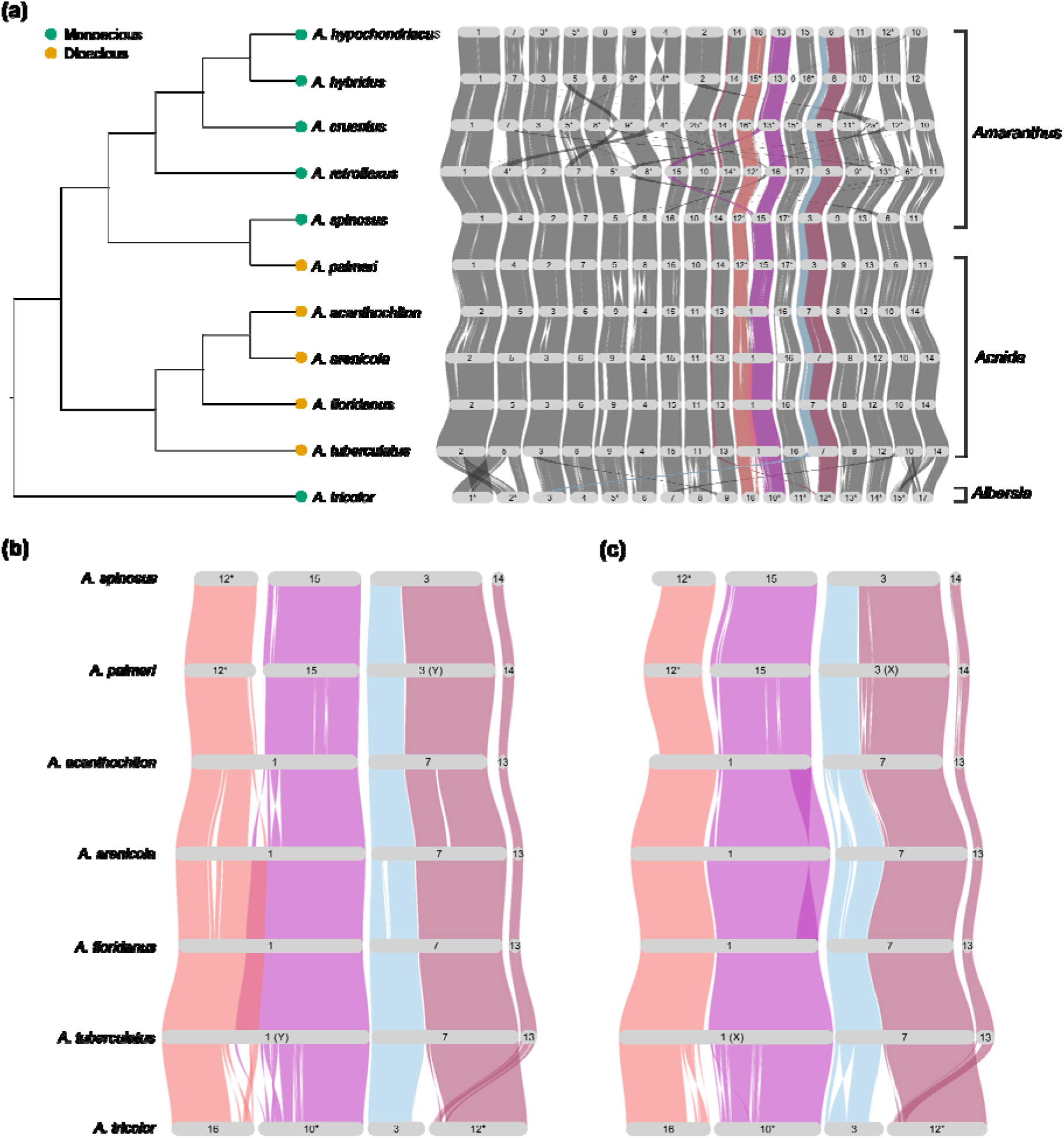
Cladogram and riparian plot of synteny among eleven *Amaranthus* species. (a) Species tree inferred using STAG and rooted with STRIDE in OrthoFinder, and syntenic relationship among the *Amaranthus* species showing the three subgenera. Hap1 used for all species with two assembled haplomes. (b,c) Syntenic block highlighting the relationship between the Y-scaffolded (Hap1) or the X-scaffolded (Hap2) chromosome of the dioecious species and homologs in the related monoecious species. For *A. arenicola* and *A. floridanus*, these designations denote the *A. tuberculatus* haplotype used for scaffolding. Asterisks depict chromosomes that were manually inverted to keep the gene order consist across the genomes.

Base-level alignment also recovered the two inversions on Chromosome 1 of *A. tuberculatus* on Chromosome 1 in *A. arenicola* and *A. floridanus*. Given that the inversions span multiple scaffolding junctions in the two species, their near-identical sizes (13.73 and 14.30 Mb; 5.24 and 6.09 Mb) reflect the shared reference, and the result is therefore consistent with conservation of the inversions in the two species. Inversions were likewise present at the same locus in *A. acanthochiton*; however, three inversions were resolved within the candidate SDR interval rather than two (Figure 4).

**FIGURE 4.**
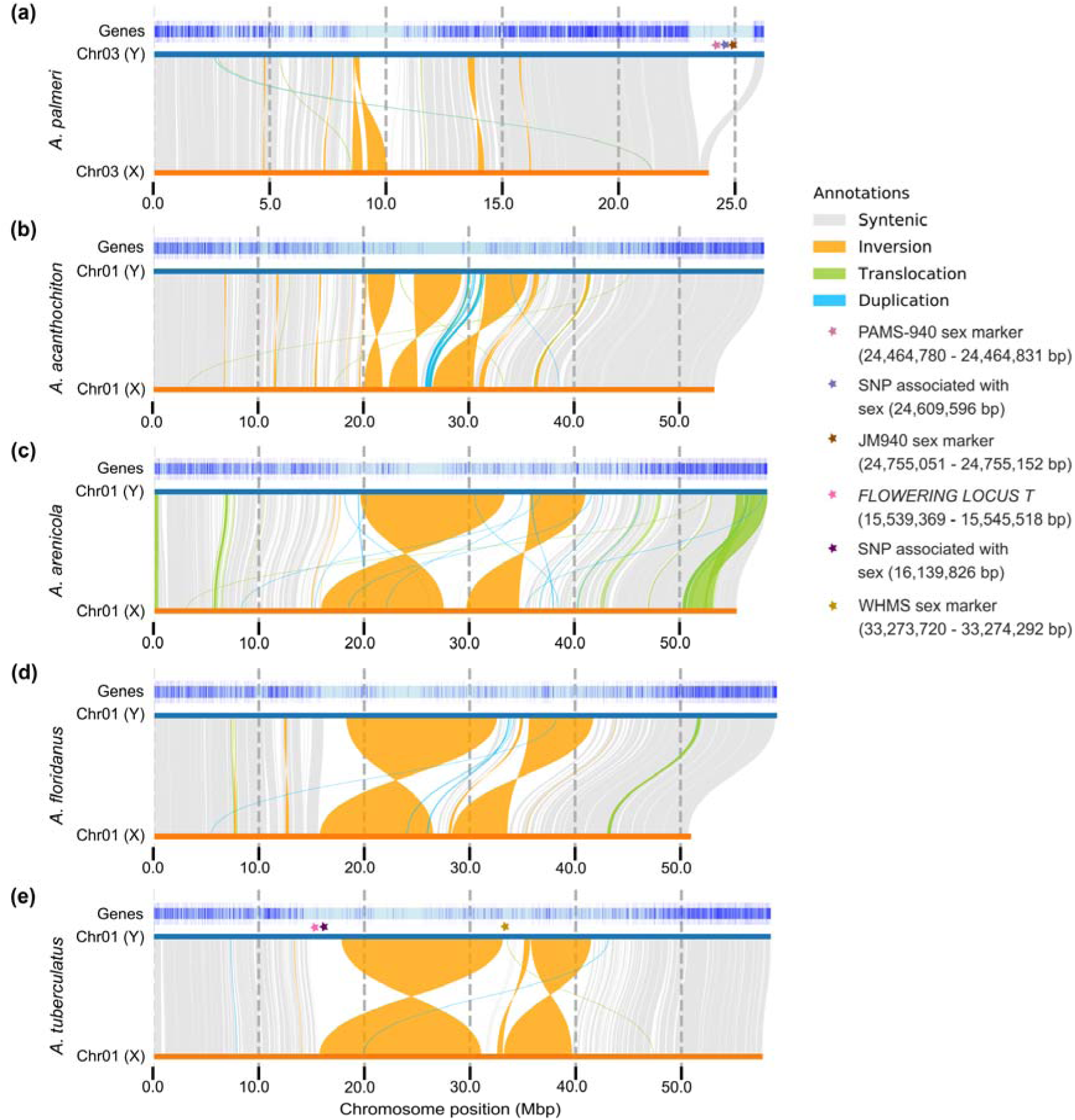
Synteny and rearrangement (SyRI) plots of Y and X comparison for a) *A. palmeri* b) *A. acanthochiton* c) *A. arenicola* d) *A. floridanus*, and e) *A. tuberculatus*. Asterisks indicate the positions of significant SNPs associated with sex in Raiyemo et al. (Raiyemo, Cutti, et al., 2025; Raiyemo, Montgomery, et al., 2025).

Comparative analysis of protein sequences, including Rf1 and TLC-domain containing protein previously reported within the candidate SDR in *A. palmeri*, to those in *A. spinosus*, a closely related monoecious species, revealed homologs on Chromosome 3 of *A. spinosus*. One monoecious copy shows only 71.6% amino acid identity across a 627-aa alignment, with 144 mismatches, and was annotated as an “unknown protein” despite containing canonical PPR motifs, indicating substantial divergence between the monoecious copy and the dioecious copy in the male-specific region (Table S8 and Figure S12). Analysis of syntenic genes among *A. acanthochiton, A. arenicola, A. floridanus*, and *A. tuberculatus* revealed 223 genes within the previously reported candidate region (Chromosome 1; 14.02–45.81 Mbp) associated with sex in *A. tuberculatus* (Raiyemo, Cutti, et al., 2025) were conserved across the species. In addition, only one allelic copy was retained arbitrarily where the assemblies carried two copies, as the GENESPACE default ploidy of one retains only the top syntenic hit at each locus. Complementary protein sequence similarity searches for genes located on Chromosome 1 identified an additional 16 genes that were conserved across species despite lacking syntenic placement, indicating conservation of gene content without conservation of genomic position (Table S9). The protein HEADING DATE 3A/FT was also conserved across the four species, although it was not contiguously assembled into the chromosome-level assembly for *A. arenicola* (Table S9).

### 3.3 Change-point analysis of synonymous divergence (K_S_)

Synonymous divergence (Ks) between X–Y gametolog pairs was examined along Chromosome 3 of *A. palmeri* and Chromosome 1 of *A. acanthochiton*, and *A. tuberculatus*, representing sex chromosome pairs in each species. After filtering out saturated estimates, datasets comprised 1,268 gametolog pairs in *A. palmeri*, 1,575 in *A. acanthochiton*, and 1,710 in *A. tuberculatus*. As a preliminary exploratory step, Generalized Additive Models (GAM) were fitted to the Ks distributions to visualize local smoothed trends along each chromosome. While smooth terms were statistically significant across all three species (*p* ≤ 0.0003), genomic position accounted for only 0.5–1.3% of Ks variance (per-species adjusted R^2^ in Figures S13 and S14), with effective degrees of freedom ranging from 5.0 to 5.8, indicating that spatial heterogeneity in synonymous divergence is detectable but weak, with no species showing the spatially discrete divergence peaks expected under a classical multi-stratum architecture.

Bayesian piecewise-constant changepoint models were further fitted to the Ks values to identify spatial shifts and discrete plateaus in synonymous divergence along the X chromosomes, allowing up to three changepoints. The models failed to converge in any of the three species (maximum R-hat = 7.1, 76.5, and 44.7 in *A. palmeri, A. acathochiton*, and *A. tuberculatus*, respectively) and simplified one-changepoint models were re-fitted. In *A. palmeri* and *A. acanthochiton*, these converged (all R-hat ≤ 1.1) but were uninformative i.e., the boundary was placed with 95% credible intervals of 3.5 – 23.7 Mbp and 0.3 – 52.4 Mbp respectively, and the two inferred segments differed negligibly in synonymous divergence (mean Ks = 0.046 and 0.037, ΔKs = 0.009 in *A. palmeri*; 0.029 and 0.027, ΔKs = 0.002 in *A. acanthochiton*). In both species the difference in expected log predictive density between the one-changepoint and zero-changepoint models fell well below the conventional threshold of four units (Δelpd = –1.5 and – 0.2), indicating no support for a stratum boundary (Figure S15 and S16, and Table S10).

In *A. tuberculatus*, the one-changepoint model identified a transition at 2.36 Mbp (95% CI: 1.82–3.04 Mbp; R-hat = 1.1 for the changepoint parameter; all other parameters R-hat = 1.0). The 1.2 Mbp credible interval is more than an order of magnitude narrower than those recovered in the other two species, and the corresponding parameter in the non-convergent three-changepoint model was recovered at a concordant position (2.27 Mbp; 95% CI: 1.83 – 3.08 Mbp), indicating that the boundary is stable across model specifications even where the additional changepoints were not identifiable. The proximal segment (0–2.36 Mbp) exhibited elevated synonymous divergence (mean Ks = 0.066) relative to the remainder of the chromosome (mean Ks = 0.050; ΔKs = 0.016). PSIS-LOO diagnostics for this model were less stable (max Pareto-k = 0.96), and the improvement over a zero-changepoint model, although exceeding four units of expected log predictive density, amounted to 1.6 standard errors (Figure S16 and Table S10). Taken together, the uniformly low synonymous divergence across all three species (mean Ks ≈ 0.03–0.07) and the general absence of resolvable divergence strata are consistent with evolutionarily young sex-determining regions, in which insufficient time has elapsed for recombination suppression to generate the stratified divergence that characterizes older sex chromosomes.

### 3.4 Gene-centric pangenome analysis

Because the four newly assembled species differ in genome size, repeat content, and chromosome structure, we sought to quantify gene family turnover across the genus and assess whether these structural differences are accompanied by variation in gene content. We therefore constructed a gene-based pangenome of *Amaranthus* species using protein-coding genes from 17 genome assemblies, spanning 11 species (*A. hypochondriacus* (Lightfoot et al., 2017), *A. cruentus* (Ma et al., 2021), *A. tricolor* (Wang et al., 2023), *A. palmeri* (Raiyemo, Montgomery, et al., 2025), *A. hybridus* (Raiyemo, Montgomery, et al., 2025), *A. retroflexus* (Raiyemo, Montgomery, et al., 2025), *A. tuberculatus* (Raiyemo, Cutti, et al., 2025), *A. spinosus, A. acanthochiton, A. arenicola*, and *A. floridanus*). The number of orthogroups (gene families) increased as more genomes were added to the analysis; however, the pangenome curve appears to be nearing a plateau at n = 16 genomes (Figure 5).

**FIGURE. 5.**
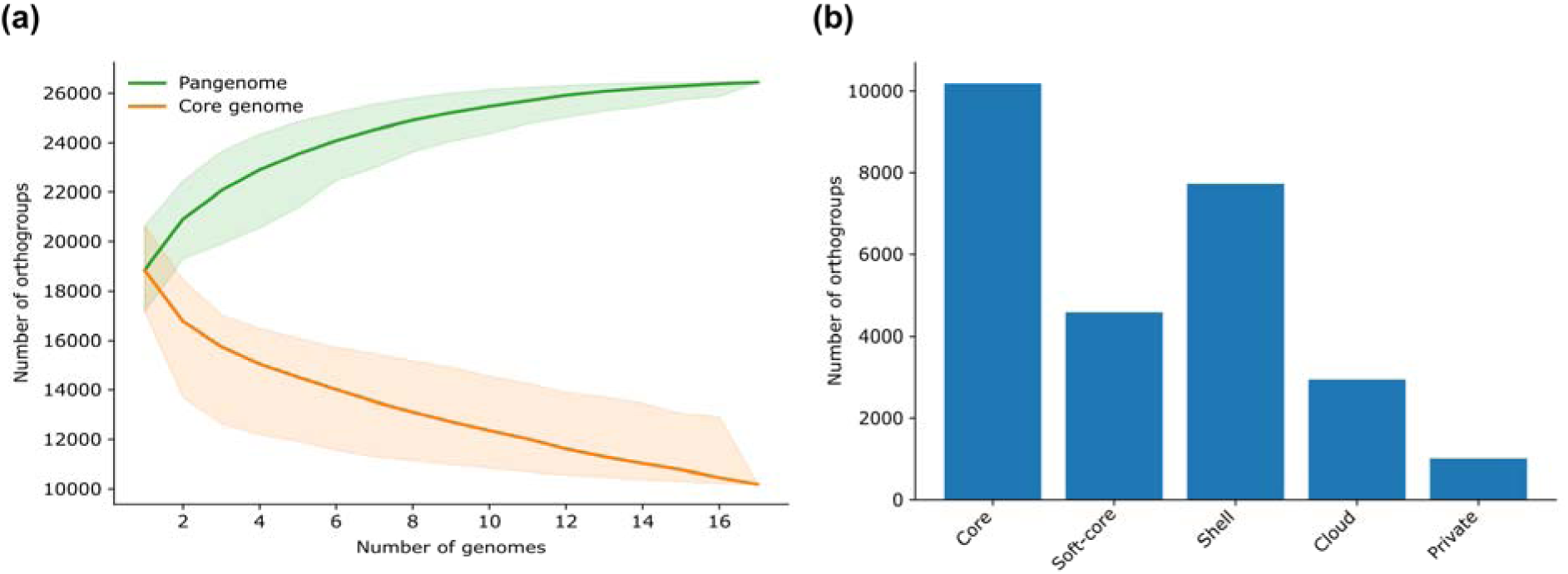
Pangenome analysis of 17 *Amaranthus* assemblies. (a) Accumulation curves of orthogroups (gene families) of the pangenome (green) and core-genome (orange). Shaded regions show the 2.5th – 97.5th percentile range across random permutations of genome order to determine how quickly the pan/core stabilizes (i.e. heterogeneity) as genomes are added. (b) bar plot of the pangenome compartments.

The pangenome consists of 26,449 orthogroups, of which 10,189 (38.5%) were core, 4,585 (17.3%) were soft-core, 7,729 (29.2%) were shell, 2,939 (11.1%) were cloud, and 1,007 (3.8%) were private (Figure 5). Similarly, after collapsing haplotypes to the species level, the pangenome consists of 11,238 (42.5%) core orthogroup, 4,692 (17.7%) soft-core orthogroup, 5,784 (21.9%) shell orthogroup, 2,493 (9.4%) cloud orthogroup, and 2,242 (8.5%) private orthogroup (Figure S17).

## 4 DISCUSSION

We report chromosome-level genome assemblies for four *Amaranthus* species, including haplotype-resolved assemblies for *A. spinosus* and *A. acanthochiton*, along with sex chromosomes of three dioecious species. Our analysis revealed the inversions and Robertsonian fusion reported for Chromosome 1 of *A. tuberculatus* (Raiyemo, Cutti, et al., 2025) were conserved in *A. acanthochiton*, which was assembled and phased *de novo*, while being consistent for the reference-scaffolded *A. arenicola* and *A. floridanus*. Together these results indicate that dioecy evolution predates speciation for the lineage. We also identified homologs of *Rf1* on Chromosome 3 of *A. spinosus*, a monoecious species that exhibits spatial separation of male and female flowers and is closely related to the dioecious *A. palmeri*. Divergence of the dioecious copy, combined with its strict male-specificity, suggests the possibility of neofunctionalization within the non-recombining male-specific Y region.

Comparative analysis of synonymous divergence across three dioecious *Amaranthus* species provided limited evidence for discrete evolutionary strata; genomic position explained at most 1.3% of Ks variance, and in *A. palmeri* and *A. acanthochiton* the one-changepoint models were statistically indistinguishable from models with no changepoint. Two features of sex determination in the genus could independently produce this outcome. First, dioecy is young. Nuclear phylogenomic dating places the *Amaranthus* crown at 2–5 Ma (Timerman et al., 2026), bounding the time available for divergence to accumulate. Second, recombination suppression appears incomplete. In *A. tuberculatus*, recombination within the sex-linked region is reduced but not eliminated, differentiation between the sexes is low, and a proportion of individuals exhibit genotype-phenotype mismatch alongside occasional leaky sex expression (Kreiner et al., 2025). Ongoing exchange between the proto–X and proto–Y would homogenize divergence regardless of the age of the system. However, the absence of strata does not necessarily follow simply from low divergence. In *Rumex hastatulus*, phased sex chromosome assemblies likewise revealed no discrete strata despite low X–Y divergence (Sacchi et al., 2024). By contrast, in the Amaranthaceae species *Spinacia tetrandra*, change–point analysis of X–Y divergence resolved two strata with median Ks values of 0.037 and 0.067 (She et al., 2025). The median gametolog divergence in *A. tuberculatus* (0.036–0.059 across segments) falls within that range, and although divergence in *A. palmeri* and *A. acanthochiton* is lower (0.033–0.035 and 0.019–0.023), no comparable stratification was detected in any of the three (medians are used for comparison because Ks distributions are right-skewed). What differs between these systems is the extent and duration of recombination suppression rather than divergence magnitude alone.

Although the one resolvable Ks transition in *A. tuberculatus* lies at ∼2.36 Mbp, this falls outside the broader SDR candidate interval (14.02–45.81 Mbp) previously delimited by structural and genome-wide association analyses (Raiyemo, Cutti, et al., 2025), and is therefore unlikely to represent stratum boundary. More generally, the weak heterogeneity we detect across the species may reflect background variation in mutation rate or local effective population size rather than recombination suppression (Jay et al., 2024; Saunders & Muyle, 2024). We note that a piecewise-constant model assumes discrete plateaus and would not detect continuous or otherwise non-stepwise variation in divergence. Taken together, the GAM and changepoint analysis indicate that Ks landscapes along *Amaranthus* sex chromosomes are largely homogeneous, providing no evidence for evolutionary strata in the species examined.

Given the gene content variation known to occur across related species, we constructed a species-level pangenome to evaluate how orthogroup presence-absence patterns reflect evolutionary processes. Across the 17 assemblies and 11 species, only 38.5% and 42.5% of orthogroups, respectively, were conserved, underscoring the extent of gene family turnover within the genus. This pattern likely reflects both genuine evolutionary divergence and methodological differences across the assemblies compared. Orthogroup assignment is sensitive to variations in gene model prediction, and the private compartment is correspondingly unevenly distributed among assemblies, with *A. cruentus* accounting for 588 or 1,007. In addition, we found that the treatment of haplotypes had a notable influence on inferred pangenome structure: analyzing haplotypes as independent genomes inflated the number of shell and cloud orthogroups, as genes present on only one haplotype were misclassified as low-frequency accessory families. Collapsing haplotypes to the species level resolved this artifact, reassigning the families to core or private categories consistent with their true species-level distribution. Our results are highly consistent with a recently constructed pangenome for the grain amaranth species complex (Ludwig et al., 2026). The apparent decrease in our strict core fraction (38.5%, or 42.5% after collapsing haplotypes) compared to their reported 74.9% is entirely driven by our expanded evolutionary scope (11 species vs. 5 species), which captures a broader, more fluid accessory genome while fundamentally validating their baseline observations. In sum, we provide high-quality genome assemblies and annotations for four *Amaranthus* species, which we utilized for comparative genomic analyses. These resources will be valuable for further studies within the genus as well as the broader Amaranthaceae family.

## AUTHOR CONTRIBUTIONS

**Damilola A. Raiyemo**: conceptualization; data curation; formal analysis; investigation, methodology; software; visualization; writing – original draft; writing – review & editing. **Isabel W. Noe**: data curation, investigation, and writing – review & editing. **Ramandeep Kaur**: formal analysis. **Lauren Whitt**: formal analysis; investigation; software; writing – original draft; writing – review & editing. **Sarah B. Carey**: formal analysis; writing – review & editing. **Haley Hale**: investigation. **Keyana J. Lewis**: investigation. **Lauren Womack**: investigation. **Alex Harkess**: conceptualization; funding acquisition; writing – review & editing. **Victor Llaca**: investigation; writing – review & editing. **Kevin Fengler**: investigation. **Eric L. Patterson**: conceptualization. **Todd A. Gaines**: conceptualization; funding acquisition; writing – review & editing. **Patrick J. Tranel**: conceptualization, funding acquisition; supervision; writing – review & editing.

## ACKNOWLEDGEMENTS

Not Applicable.

## DATA AVAILABILITY STATEMENT

Raw reads data from this study are available through the National Center for Biotechnology Information (NCBI) under project numbers PRJNA1463991, PRJNA1464092, PRJNA1463989, PRJNA1464093, and PRJNA1463199. This Whole Genome Shotgun project has been deposited at DDBJ/ENA/GenBank under the accessions JCAAIL000000000, JCAAIM000000000, JCAAIN000000000, JCAAIO000000000, JCASDJ000000000, JCASDK000000000, JCASDL000000000, and JCASDM000000000. Codes used for bioinformatics analyses in this project are available on GitHub (https://github.com/Alexdami17/Amaranthus-genome-assembly).

## SUPPLEMENTAL MATERIAL STATEMENT

Figure S1. Whole-genome pairwise self-alignment of a) *A. spinosus* Hap1 and b) *A. spinosus* Hap2.

Figure S2. Whole-genome pairwise alignment between a) *A. palmeri* Hap1 and *A. spinosus* Hap1, and b) *A. palmeri* Hap1 and *A. spinosus* Hap2.

Figure S3. Whole-genome pairwise self-alignment of a) *A. acanthochiton* Hap1, and b) *A. acanthochiton* Hap2.

Figure S4. Whole-genome pairwise alignment between a) *A. acanthochiton* Hap1 and *A. acanthochiton* Hap2, and b) *A. tuberculatus* Hap1 and *A. acanthochiton* Hap1.

Figure S5. Whole-genome pairwise (a) self-alignment of *A. arenicola* Hap1, and (b) alignment between *A. tuberculatus* Hap1 and *A. arenicola* Hap1.

Figure S6. Whole-genome pairwise (a) self-alignment of *A. arenicola* Hap2, and (b) alignment between *A. tuberculatus* Hap2 and *A. arenicola* Hap2.

Figure S7. Whole-genome pairwise alignment between *A. arenicola* Hap1 and *A. arenicola* Hap2.

Figure S8. Whole-genome pairwise (a) self-alignment of *A. floridanus* Hap1, and (b) alignment between *A. tuberculatus* Hap1 and *A. floridanus* Hap1.

Figure S9. Whole-genome pairwise (a) self-alignment of *A. floridanus* Hap2, and (b) alignment between *A. tuberculatus* Hap2 and *A. floridanus* Hap2.

Figure S10. Whole-genome pairwise alignment between *A. floridanus* Hap1 and *A. floridanus* Hap2.

Figure S11. Synteny and rearrangement plot (SyRI) between *A. palmeri* Hap1 and *A. spinosus* Hap1, providing a structural variant-level view of the same comparison shown as a whole-genome dotplot in Figure S2a.

Figure S12. Phylogenetic tree of Rf1 protein sequences in *A. palmeri* and its homologs in other *Amaranthus* species (*A. hypochondriacus, A. hybridus, A. cruentus, A. retroflexus, A. spinosus, A. acanhtochiton, A. arenicola, A. floridanus, A. tuberculatus*, and *A. tricolor*).

Figure S13. Synonymous divergence (Ks) between one-to-one X–Y gametologs along the X chromosome (Chr03) of *A. palmeri*.

Figure S14. Synonymous divergence (Ks) between one-to-one X–Y gametologs along the X chromosome (Chr01) of *A. acanthochiton* (a), *A. arenicola* (b), *A. floridanus* (c), and *A. tuberculatus* (d).

Figure S15. Distribution of synonymous divergence (Ks) values from one-to-one orthologs along X chromosome (Chr03) of *A. palmeri*.

Figure S16. Distribution of synonymous divergence (Ks) values from one-to-one orthologs along the X chromosome (Chr01) of *A. acanthochiton* (a), *A. arenicola* (b), *A. floridanus* (c), and *A. tuberculatus* (d).

Figure S17. Pangenome analysis of 11 *Amaranthus* species genomes.

## Supplemental Tables S1-S10

Table S1. *Amaranthus* species with available genome assemblies.

Table S2. Read length and estimated coverage for species sequenced and assembled in this study.

Table S3. Repeat composition of the genome assemblies.

Table S4. Summary of BLAST query of telomeric repeat against the genome assemblies.

Table S5. Inversions between the two haplotype assemblies of *Amaranthus acanthochiton*.

Table S6. Inversions between the two haplotype assemblies of *Amaranthus arenicola*.

Table S7. Inversions between the two haplotype assemblies of *Amaranthus floridanus*.

Table S8. MMseqs easy-search of the genes within the male-specific region of *A. palmeri* in monoecious *A. spinosus*.

Table S9. Genes conserved by synteny and sequence similarity across four *Amaranthus* species.

Table S10. Bayesian changepoint parameter estimates.

## Abbreviations

BUSCO: benchmarking universal single-copy orthologs;
SDR: sex-determining region;
MSY: male-specific Y;
GRIN: germplasm resources information network;
HMW: high molecular weight;
LTR: long terminal repeat;
LAI: long-terminal-repeat assembly index;
LINE: long interspersed nuclear element;
TE: transposable element;
tRNA: transfer RNA;
BLASTN: basic local alignment search tool for nucleotides;
CDS: coding sequence;
UTRs: untranslated regions;
GAM: generalized additive model;
MCMC: Markov Chain Monte Carlo.

## REFERENCES

Adhikary, D., & Pratt, D. B. (2015). Morphologic and taxonomic analysis of the weedy and cultivated *Amaranthus hybridus* species complex. Systematic Botany, 40(2), 604–610. 10.1600/036364415X688376

Alonge, M., Lebeigle, L., Kirsche, M., Jenike, K., Ou, S., Aganezov, S., Wang, X., Lippman, Z. B., Schatz, M. C., & Soyk, S. (2022). Automated assembly scaffolding using RagTag elevates a new tomato system for high-throughput genome editing. Genome Biology, 23(1). 10.1186/s13059-022-02823-7

Astashyn, A., Tvedte, E. S., Sweeney, D., Sapojnikov, V., Bouk, N., Joukov, V., Mozes, E., Strope, P. K., Sylla, P. M., Wagner, L., Bidwell, S. L., Brown, L. C., Clark, K., Davis, E. W., Smith-White, B., Hlavina, W., Pruitt, K. D., Schneider, V. A., & Murphy, T. D. (2024). Rapid and sensitive detection of genome contamination at scale with FCS-GX. Genome Biology, 25(1). 10.1186/s13059-024-03198-7

Bayón, N. D. (2022). Identifying the weedy amaranths (Amaranthus, Amaranthaceae) of South America. Advances in Weed Science, 40(spe2), 1–9. 10.51694/advweedsci/2022;40:amaranthus007

Bushnell, B. (2014). BBTools software package. https://sourceforge.net/projects/bbmap

Cabanettes, F., & Klopp, C. (2018). D-GENIES: Dot plot large genomes in an interactive, efficient and simple way. PeerJ, 2018(6). 10.7717/peerj.4958

Camacho, C., Coulouris, G., Avagyan, V., Ma, N., Papadopoulos, J., Bealer, K., & Madden, T. L. (2009). BLAST+: Architecture and applications. BMC Bioinformatics, 10, 1–9. 10.1186/1471-2105-10-421

Capella-Gutiérrez, S., Silla-Martínez, J. M., & Gabaldón, T. (2009). trimAl: A tool for automated alignment trimming in large-scale phylogenetic analyses. Bioinformatics, 25(15), 1972–1973. 10.1093/bioinformatics/btp348

Carey, S. B., Aközbek, L., Lovell, J. T., Jenkins, J., Healey, A. L., Shu, S., Grabowski, P., Yocca, A., Stewart, A., Jones, T., Barry, K., Rajasekar, S., Talag, J., Scutt, C., Lowry, P. P., Munzinger, J., Knox, E. B., Soltis, D. E., Soltis, P. S., … Harkess, A. (2024). ZW sex chromosome structure in *Amborella trichopoda*. Nature Plants, 10(12), 1944–1954. 10.1038/s41477-024-01858-x

Chan, P. P., Lin, B. Y., Mak, A. J., & Lowe, T. M. (2021). TRNAscan-SE 2.0: Improved detection and functional classification of transfer RNA genes. Nucleic Acids Research, 49(16), 9077–9096. 10.1093/nar/gkab688

Chauhan, B. S., & Abugho, S. B. (2012). Phenotypic plasticity of spiny amaranth (*Amaranthus spinosus*) and longfruited primrose-willow (*Ludwigia octovalvis*) in response to rice interference. Weed Science, 60(3), 411–415. 10.1614/ws-d-11-00158.1

Chauhan, B. S., & Johnson, D. E. (2009). Germination ecology of spiny (*Amaranthus spinosus*) and slender amaranth (*A. viridis*): Troublesome weeds of direct-seeded rice. Weed Science, 57(4), 379–385. 10.1614/ws-08-179.1

Cheng, H., Concepcion, G. T., Feng, X., Zhang, H., & Li, H. (2021). Haplotype-resolved de novo assembly using phased assembly graphs with hifiasm. Nature Methods, 18(2), 170–175. 10.1038/s41592-020-01056-5

Dainat, J. (2022). Another Gtf/Gff analysis toolkit (AGAT): Resolve interoperability issues and accomplish more with your annotations. Plant and Animal Genome XXIX Conference. https://github.com/NBISweden/AGAT

Danecek, P., Bonfield, J. K., Liddle, J., Marshall, J., Ohan, V., Pollard, M. O., Whitwham, A., Keane, T., McCarthy, S. A., & Davies, R. M. (2021). Twelve years of SAMtools and BCFtools. GigaScience, 10(2), 1–4. 10.1093/gigascience/giab008

De Coster, W., & Rademakers, R. (2023). NanoPack2: population-scale evaluation of long-read sequencing data. Bioinformatics, 39(5). 10.1093/bioinformatics/btad311

Dudchenko, O., Shamim, M. S., Batra, S. S., Durand, N. C., Musial, N. T., Mostofa, R., Pham, M., Glenn St Hilaire, B., Yao, W., Stamenova, E., Hoeger, M., Nyquist, S. K., Korchina, V., Pletch, K., Flanagan, J. P., Tomaszewicz, A., McAloose, D., Pérez Estrada, C., Novak, B. J., … Aiden, E. L. (2018). The Juicebox Assembly Tools module facilitates de novo assembly of mammalian genomes with chromosome-length scaffolds for under $1000. *BioRxiv*. 10.1101/254797

Durand, N. C., Robinson, J. T., Shamim, M. S., Machol, I., Mesirov, J. P., Lander, E. S., & Aiden, E. L. (2016). Juicebox provides a visualization system for Hi-C contact maps with unlimited zoom. Cell Systems, 3(1), 99–101. 10.1016/j.cels.2015.07.012

Eliasson, U. H. (1988). Floral morphology and taxonomic relations among the genera of Amaranthaceae in the New World and the Hawaiian Islands. Botanical Journal of the Linnean Society, 96(3), 235–283. 10.1111/j.1095-8339.1988.tb00683.x

Emms, D. M., & Kelly, S. (2019). OrthoFinder: Phylogenetic orthology inference for comparative genomics. Genome Biology, 20(1). 10.1186/s13059-019-1832-y

Faccenda, K., & Ross, M. C. (2024). New naturalization records for *Amaranthus* in the Hawaiian Islands. Bishop Museum Occasional Papers, (156), 23–32.

Flynn, J. M., Hubley, R., Goubert, C., Rosen, J., Clark, A. G., Feschotte, C., & Smit, A. F. (2020). RepeatModeler2 for automated genomic discovery of transposable element families. Proceedings of the National Academy of Sciences of the United States of America, 117(17), 9451–9457. 10.1073/pnas.1921046117

Gabriel, L., Brůna, T., Hoff, K. J., Ebel, M., Lomsadze, A., Borodovsky, M., & Stanke, M. (2024). BRAKER3: Fully automated genome annotation using RNA-seq and protein evidence with GeneMark-ETP, AUGUSTUS, and TSEBRA. Genome Research, 34(5), 769–777. 10.1101/gr.278090.123

Goel, M., & Schneeberger, K. (2022). plotsr: Visualizing structural similarities and rearrangements between multiple genomes. Bioinformatics, 38(10), 2922–2926. 10.1093/bioinformatics/btac196

Goel, M., Sun, H., Jiao, W. B., & Schneeberger, K. (2019). SyRI: finding genomic rearrangements and local sequence differences from whole-genome assemblies. Genome Biology, 20(1), 1–13. 10.1186/s13059-019-1911-0

Grant, W. F. (1959). Cytogenetic studies in *Amaranthus*. Canadian Journal of Botany, 37, 413–417. 10.1139/b59-032

Hart, A. J., Ginzburg, S., Xu, M., Fisher, C. R., Rahmatpour, N., Mitton, J. B., Paul, R., & Wegrzyn, J. L. (2020). EnTAP: Bringing faster and smarter functional annotation to non-model eukaryotic transcriptomes. Molecular Ecology Resources, 20(2), 591–604. 10.1111/1755-0998.13106

Huang, N., & Li, H. (2023). compleasm: a faster and more accurate reimplementation of BUSCO. Bioinformatics, 39(10). 10.1093/bioinformatics/btad595

Iamonico, D. (2020). Nomenclatural survey of the genus Amaranthus (Amaranthaceae). 11. dioecious Amaranthus species belonging to the sect. Saueranthus. Darwiniana, 8(2), 567–575. 10.14522/darwiniana.2020.82.898

Jay, P., Jeffries, D., Hartmann, F. E., Véber, A., & Giraud, T. (2024). Why do sex chromosomes progressively lose recombination? Trends in Genetics, 40(7), 564–579. 10.1016/j.tig.2024.03.005

Katoh, K., & Standley, D. M. (2013). MAFFT multiple sequence alignment software version 7: Improvements in performance and usability. Molecular Biology and Evolution, 30(4), 772–780. 10.1093/molbev/mst010

Kaur, H., Shannon, L. M., & Samac, D. A. (2024). A stepwise guide for pangenome development in crop plants: an alfalfa (*Medicago sativa*) case study. BMC Genomics, 25(1), 1022. 10.1186/s12864-024-10931-w

Kreiner, J. M., Montgomery, J. S., Todesco, M., Bercovich, N., Gong, Y., Elphinstone, C., Tranel, P. J., Rieseberg, L. H., & Wright, S. I. (2025). The evolution of separate sexes in waterhemp is associated with surprising chromosomal diversity and complexity. PLOS Biology, 23(6). 10.1371/journal.pbio.3003254

Li, H. (2013). Aligning sequence reads, clone sequences and assembly contigs with BWA-MEM. *ArXiv:1303.3997v2, 00*(00), 1–3. http://arxiv.org/abs/1303.3997

Li, H. (2018). Minimap2: Pairwise alignment for nucleotide sequences. Bioinformatics, 34(18), 3094–3100. 10.1093/bioinformatics/bty191

Li, H. (2023). Protein-to-genome alignment with miniprot. Bioinformatics, 39(1). 10.1093/bioinformatics/btad014

Lightfoot, D. J., Jarvis, D. E., Ramaraj, T., Lee, R., Jellen, E. N., & Maughan, P. J. (2017). Single-molecule sequencing and Hi-C-based proximity-guided assembly of amaranth (*Amaranthus hypochondriacus*) chromosomes provide insights into genome evolution. BMC Biology, 15(1), 1–15. 10.1186/s12915-017-0412-4

Lovell, J. T., Sreedasyam, A., Schranz, M. E., Wilson, M., Carlson, J. W., Harkess, A., Emms, D., Goodstein, D. M., & Schmutz, J. (2022). GENESPACE tracks regions of interest and gene copy number variation across multiple genomes. ELife, 11, 1–20. 10.7554/ELIFE.78526

Ludwig, E., Winkler, T. S., & Stetter, M. G. (2026). The grain amaranth pangenome reveals domestication-associated changes in diversity and function of structural variation. bioRxiv. 10.64898/2026.01.08.698315

Ma, X., Vaistij, F. E., Li, Y., Jansen van Rensburg, W. S., Harvey, S., Bairu, M. W., Venter, S. L., Mavengahama, S., Ning, Z., Graham, I. A., Van Deynze, A., Van de Peer, Y., & Denby, K. J. (2021). A chromosome level *Amaranthus cruentus* genome assembly highlights gene family evolution and biosynthetic gene clusters that may underpin the nutritional value of this traditional crop. The Plant Journal, 107(2), 613–628. 10.1111/tpj.15298

Mallory, M. A., Hall, R. V., McNabb, A. R., Pratt, D. B., Jellen, E. N., & Maughan, P. J. (2008). Development and characterization of microsatellite markers for the grain amaranths. Crop Science, 48(3), 1098–1106. 10.2135/cropsci2007.08.0457

Minnis, P. E. (1991). Famine foods of the Northern American desert borderlands in historical context. Journal of Ethnobiology, 11(2), 231–257.

Montgomery, J. S., Giacomini, D. A., Weigel, D., & Tranel, P. J. (2021). Male-specific Y-chromosomal regions in waterhemp (*Amaranthus tuberculatus*) and Palmer amaranth (*Amaranthus palmeri*). New Phytologist, 229(6), 3522–3533. 10.1111/nph.17108

Montgomery, J. S., Sadeque, A., Giacomini, D. A., Brown, P. J., & Tranel, P. J. (2019). Sex-specific markers for waterhemp (*Amaranthus tuberculatus*) and Palmer amaranth (*Amaranthus palmeri*). Weed Science, 67(4), 412–418. 10.1017/wsc.2019.27

Nandula, V. K., Wright, A. A., Bond, J. A., Ray, J. D., Eubank, T. W., & Molin, W. T. (2014). EPSPS amplification in glyphosate-resistant spiny amaranth (*Amaranthus spinosus*): A case of gene transfer via interspecific hybridization from glyphosate-resistant Palmer amaranth (*Amaranthus palmeri*). Pest Management Science, 70(12), 1902–1909. 10.1002/ps.3754

Odero, D. C., & Wright, A. L. (2022). Preemergence and postemergence spiny amaranth (*Amaranthus spinosus*) and common lambsquarters (*Chenopodium album*) control in lettuce on organic soils. Weed Technology, 36(4), 531–536. 10.1017/wet.2022.41

Open2C, Abdennur, N., Fudenberg, G., Flyamer, I. M., Galitsyna, A. A., Goloborodko, A., Imakaev, M., & Venev, S. V. (2024). Pairtools: From sequencing data to chromosome contacts. PLoS Computational Biology, 20(5 May). 10.1371/journal.pcbi.1012164

Ou, S., & Jiang, N. (2018). LTR_retriever: A highly accurate and sensitive program for identification of long terminal repeat retrotransposons. Plant Physiology, 176(2), 1410–1422. 10.1104/pp.17.01310

Raiyemo, D. A., Bobadilla, L. K., & Tranel, P. J. (2023). Genomic profiling of dioecious *Amaranthus* species provides novel insights into species relatedness and sex genes. BMC Biology, 21(37), 1–18. 10.1186/s12915-023-01539-9

Raiyemo, D. A., Cutti, L., Patterson, E. L., Llaca, V., Fengler, K., Montgomery, J. S., Morran, S., Gaines, T. A., & Tranel, P. J. (2025). Analyses of contiguous reference genomes of *Amaranthus tuberculatus* highlight the landscape of the sex-associated region and PEBP gene family diversity. BMC Genomics, 26(1), 988. 10.1186/s12864-025-12181-w

Raiyemo, D. A., Montgomery, J. S., Cutti, L., Abdollahi, F., Llaca, V., Fengler, K., Lopez, A. J., Morran, S., Saski, C. A., Nelson, D. R., Patterson, E. L., Gaines, T. A., & Tranel, P. J. (2025). Chromosome-level assemblies of *Amaranthus palmeri, Amaranthus retroflexus*, and *Amaranthus hybridus* allow for genomic comparisons and identification of a sex-determining region. Plant Journal, 121(4), 1–17. 10.1111/tpj.70027

Raiyemo, D. A., & Tranel, P. J. (2023). Comparative analysis of dioecious *Amaranthus* plastomes and phylogenomic implications within Amaranthaceae s.s. BMC Ecology and Evolution, 23(1), 15. 10.1186/s12862-023-02121-1

Rhie, A., Walenz, B. P., Koren, S., & Phillippy, A. M. (2020). Merqury: Reference-free quality, completeness, and phasing assessment for genome assemblies. Genome Biology, 21(1). 10.1186/s13059-020-02134-9

Sacchi, B., Humphries, Z., Kružlicová, J., Bodláková, M., Pyne, C., Choudhury, B. I., Gong, Y., Bačovský, V., Hobza, R., Barrett, S. C. H., & Wright, S. I. (2024). Phased assembly of neo-sex chromosomes reveals extensive Y degeneration and rapid genome evolution in *Rumex hastatulus*. Molecular Biology and Evolution, 41(4). 10.1093/molbev/msae074

Sarker, U., & Oba, S. (2019). Nutraceuticals, antioxidant pigments, and phytochemicals in the leaves of *Amaranthus spinosus* and *Amaranthus viridis* weedy species. Scientific Reports, 9(1). 10.1038/s41598-019-50977-5

Sauer, J. D. (1955). Revision of the dioecious amaranths. Madroño, 13(1), 5–46.

Sauer, J. D. (1957). Recent migration and evolution of the dioecious amaranths. Evolution, 11(1), 11–31.

Sauer, J. D. (1972). The dioecious amaranths: A new species name and major range extensions. Madrono, 21(6), 426.

Sauer, J. D. (1967). The grain amaranths and their relatives: A revised taxonomic and geographic survey. Annals of the Missouri Botanical Garden, 54(2), 103–137.

Saunders, P. A., & Muyle, A. (2024). Sex chromosome evolution: Hallmarks and question marks. Molecular Biology and Evolution, 41(11). 10.1093/molbev/msae218

She, H., Liu, Z., Xu, Z., Zhang, H., Wu, J., Wang, X., Cheng, F., Charlesworth, D., & Qian, W. (2025). Genome sequence of the wild species, *Spinacia tetrandra*, including a phased sequence of the extensive sex-linked region, revealing partial degeneration in evolutionary strata with unusual properties. New Phytologist, 246(6), 2765–2781. 10.1111/nph.70165

Sim, S. B., Corpuz, R. L., Simmonds, T. J., & Geib, S. M. (2022). HiFiAdapterFilt, a memory efficient read processing pipeline, prevents occurrence of adapter sequence in PacBio HiFi reads and their negative impacts on genome assembly. BMC Genomics, 23(1). 10.1186/s12864-022-08375-1

Simão, F. A., Waterhouse, R. M., Ioannidis, P., Kriventseva, E. V., & Zdobnov, E. M. (2015). BUSCO: Assessing genome assembly and annotation completeness with single-copy orthologs. Bioinformatics, 31(19), 3210–3212. 10.1093/bioinformatics/btv351

Steinegger, M., & Söding, J. (2017). MMseqs2 enables sensitive protein sequence searching for the analysis of massive data sets. Nature Biotechnology, 35(11), 1026–1028. 10.1038/nbt.3988

Stetter, M. G., & Schmid, K. J. (2017). Analysis of phylogenetic relationships and genome size evolution of the *Amaranthus* genus using GBS indicates the ancestors of an ancient crop. Molecular Phylogenetics and Evolution, 109, 80–92. 10.1016/j.ympev.2016.12.029

Suyama, M., Torrents, D., & Bork, P. (2006). PAL2NAL: Robust conversion of protein sequence alignments into the corresponding codon alignments. Nucleic Acids Research, 34(W1), W609–W612. 10.1093/nar/gkl315

Tegenfeldt, F., Kuznetsov, D., Manni, M., Berkeley, M., Zdobnov, E. M., & Kriventseva, E. V. (2025). OrthoDB and BUSCO update: annotation of orthologs with wider sampling of genomes. Nucleic Acids Research, 53(D1), D516–D522. 10.1093/nar/gkae987

Tettelin, H., & Medini, D. (Eds.). (2020). The Pangenome: Diversity, Dynamics and Evolution of Genomes. Springer Cham. 10.1007/978-3-030-38281-0

Timerman, D., Leung, J., & Eaton, D. A. R. (2026). Organelle Capture, Lineage-Specific Genomic Responses, and the Lability of Dioecy in Amaranthus. bioRxiv. 10.64898/2026.06.13.732039

Trucco, F., Jeschke, M. R., Rayburn, A. L., & Tranel, P. J. (2005). Promiscuity in weedy amaranths: high frequency of female tall waterhemp (*Amaranthus tuberculatus*) × smooth pigweed (*A. hybridus*) hybridization under field conditions. Weed Science, 53(1), 46–54. 10.1614/ws-04-103r

Vaser, R., Sović, I., Nagarajan, N., & Šikić, M. (2017). Fast and accurate de novo genome assembly from long uncorrected reads. Genome Research, 27(5), 737–746. 10.1101/gr.214270.116

Vehtari, A., Gelman, A., & Gabry, J. (2017). Practical Bayesian model evaluation using leave-one-out cross-validation and WAIC. Statistics and Computing, 27(5), 1413–1432. 10.1007/s11222-016-9696-4

Wang, H., Xu, D., Wang, S., Wang, A., Lei, L., Jiang, F., Yang, B., Yuan, L., Chen, R., Zhang, Y., & Fan, W. (2023). Chromosome-scale *Amaranthus tricolor* genome provides insights into the evolution of the genus *Amaranthus* and the mechanism of betalain biosynthesis. DNA Research, 30(1), 1–15. 10.1093/dnares/dsac050

Waselkov, K. E., Boleda, A. S., & Olsen, K. M. (2018). A phylogeny of the genus *Amaranthus* (Amaranthaceae) based on several low-copy nuclear loci and chloroplast regions. Systematic Botany, 43(2), 439–458. 10.1600/036364418X697193

Wong, T. K., Ly-Trong, N., Ren, H., Baños, H., Roger, A. J., Susko, E., Bielow, C., De Maio, N., Goldman, N., Hahn, M. W., Huttley, G., Lanfear, R., & Quang Minh, B. (2026). IQ-TREE 3: Phylogenomic Inference Software using Complex Evolutionary Models. Molecular Biology and Evolution, 43(5), msag117. 10.1093/molbev/msag117

Xu, H., Xiang, N., Du, W., Zhang, J., & Zhang, Y. (2022). Genetic variation and structure of complete chloroplast genome in alien monoecious and dioecious *Amaranthus* weeds. Scientific Reports, 12(1), 1–9. 10.1038/s41598-022-11983-2

Zhang, Z. (2022). KaKs_Calculator 3.0: Calculating selective pressure on coding and non-coding sequences. *Genomics*, Proteomics and Bioinformatics, 20(3), 536–540. 10.1016/j.gpb.2021.12.002

Zhou, C., Brown, M., Blaxter, M., McCarthy, S. A., & Durbin, R. (2025). Oatk: a de novo assembly tool for complex plant organelle genomes. Genome Biology, 26(1), 235. 10.1186/s13059-025-03676-6

Zhou, C., McCarthy, S. A., & Durbin, R. (2023). YaHS: yet another Hi-C scaffolding tool. Bioinformatics, 39(1), btac808. 10.1093/bioinformatics/btac808

