## Supplementary figures for "Chromosome-level genome assemblies and annotations of Amaranthus spinosus, Amaranthus acanthochiton, Amaranthus arenicola, and Amaranthus floridanus"

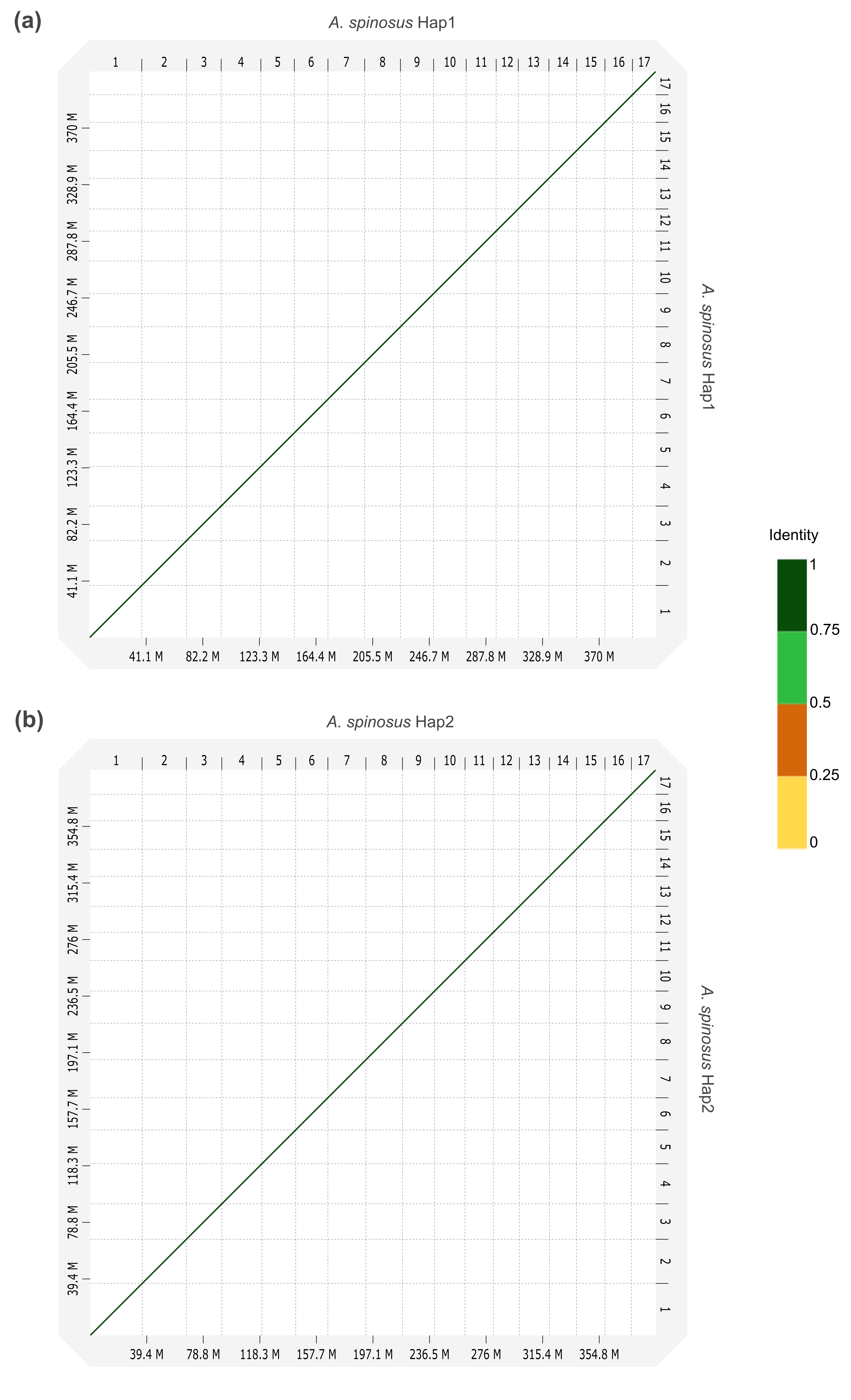

**Figure S1.** Whole-genome pairwise self-alignment of (a) *A. spinosus* Hap1, and (b) *A. spinosus* Hap2.

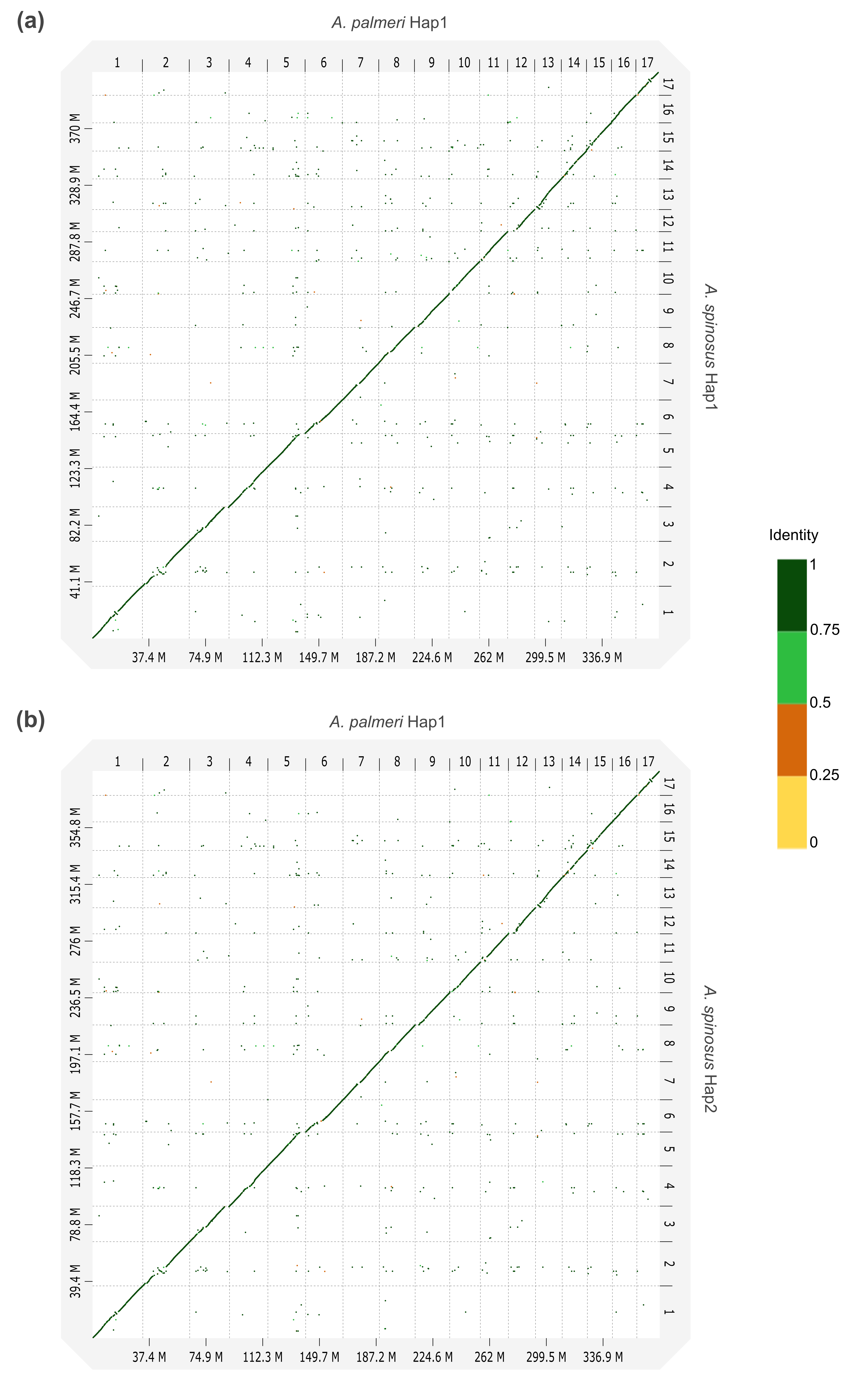

**Figure S2.** Whole-genome pairwise alignment between (a) *A. palmeri* Hap1 and *A. spinosus* Hap1, and (b) *A. palmeri* Hap1 and *A. spinosus* Hap2.

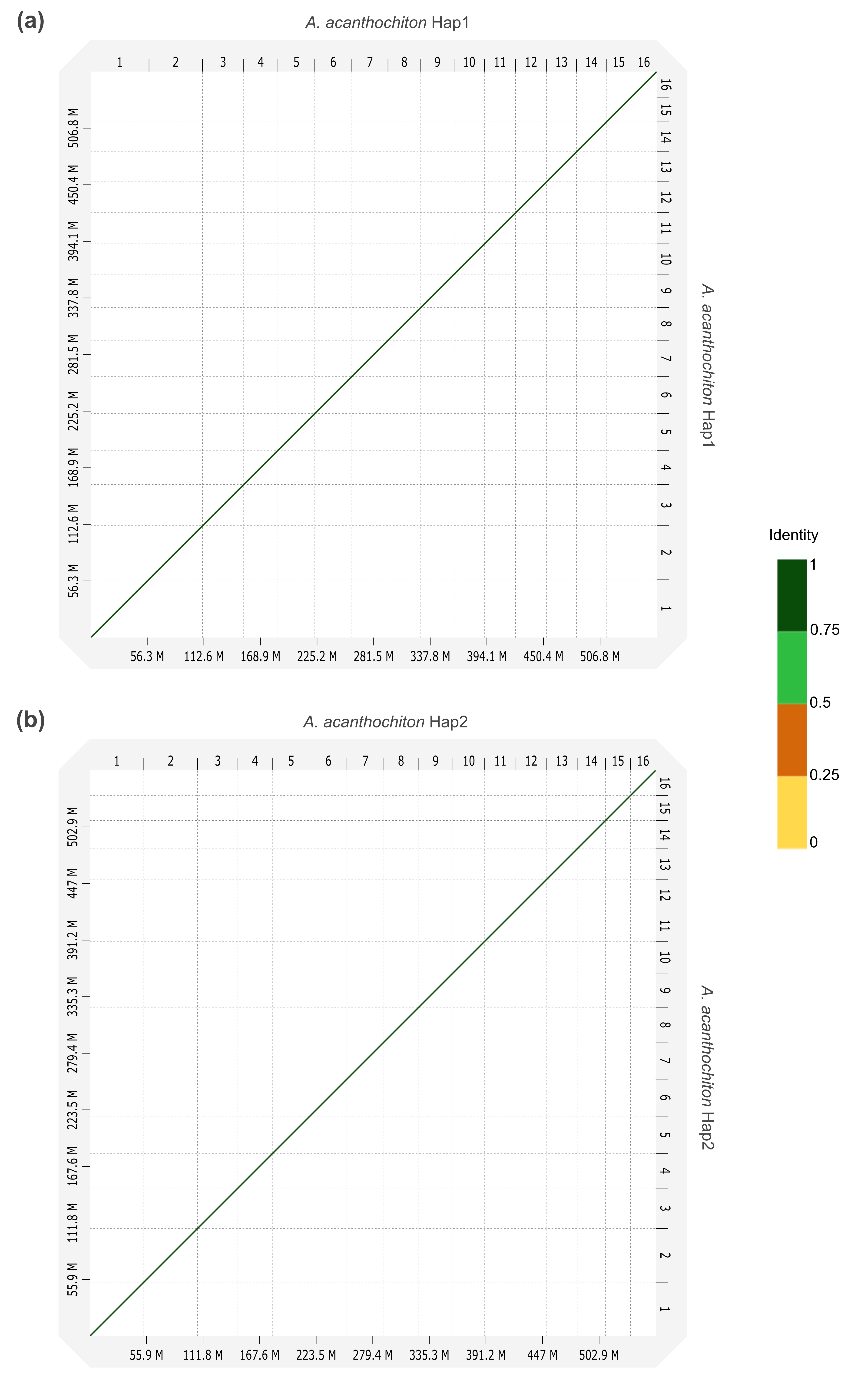

**Figure S3.** Whole-genome pairwise self-alignment of (a) *A. acanthochiton* Hap1, and (b) *A. acanthochiton* Hap2.

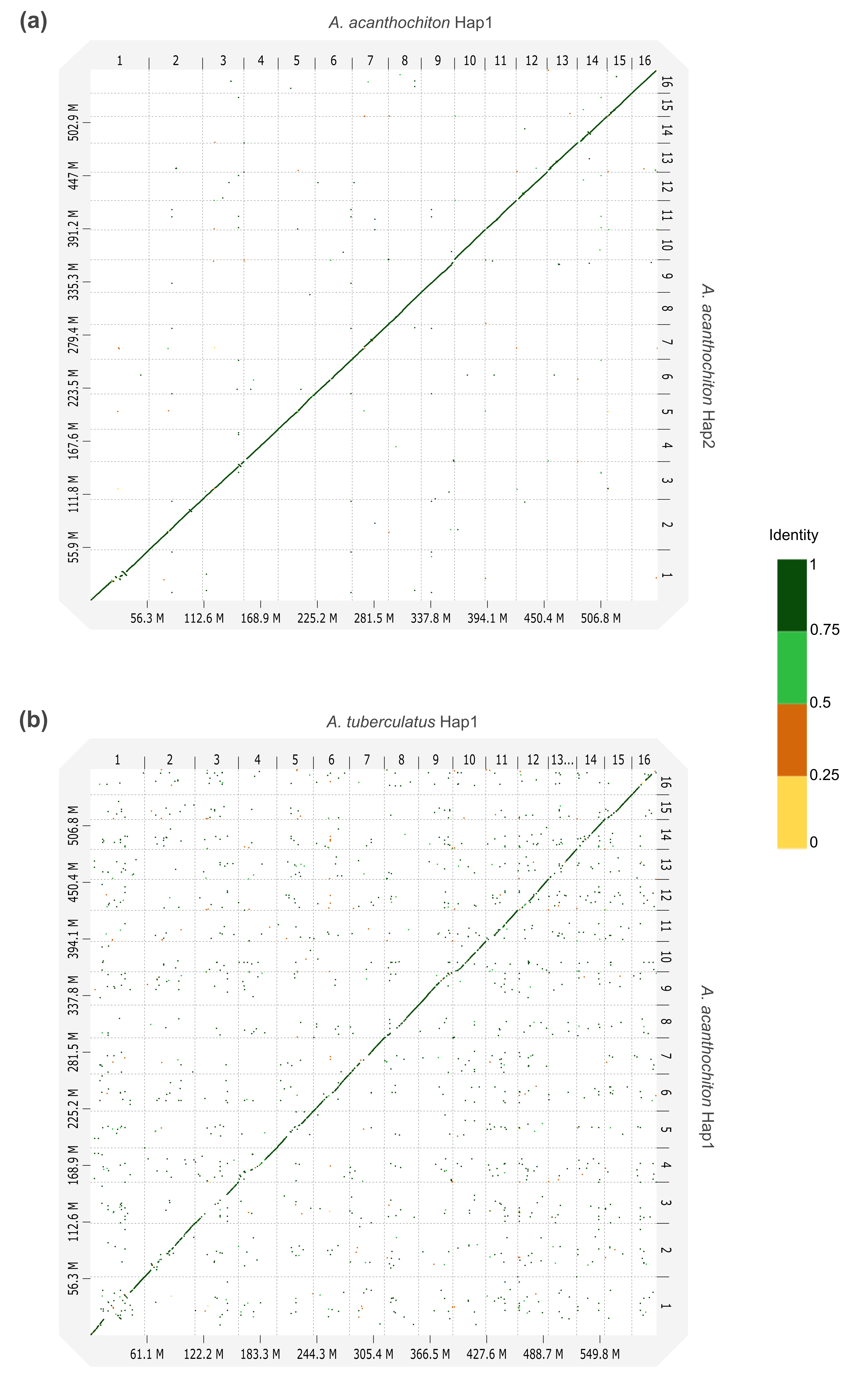

**Figure S4.** Whole-genome pairwise alignment between (a) *A. acanthochiton* Hap1 and *A. acanthochiton* Hap2, and (b) *A. tuberculatus* Hap1 and *A. acanthochiton* Hap1.

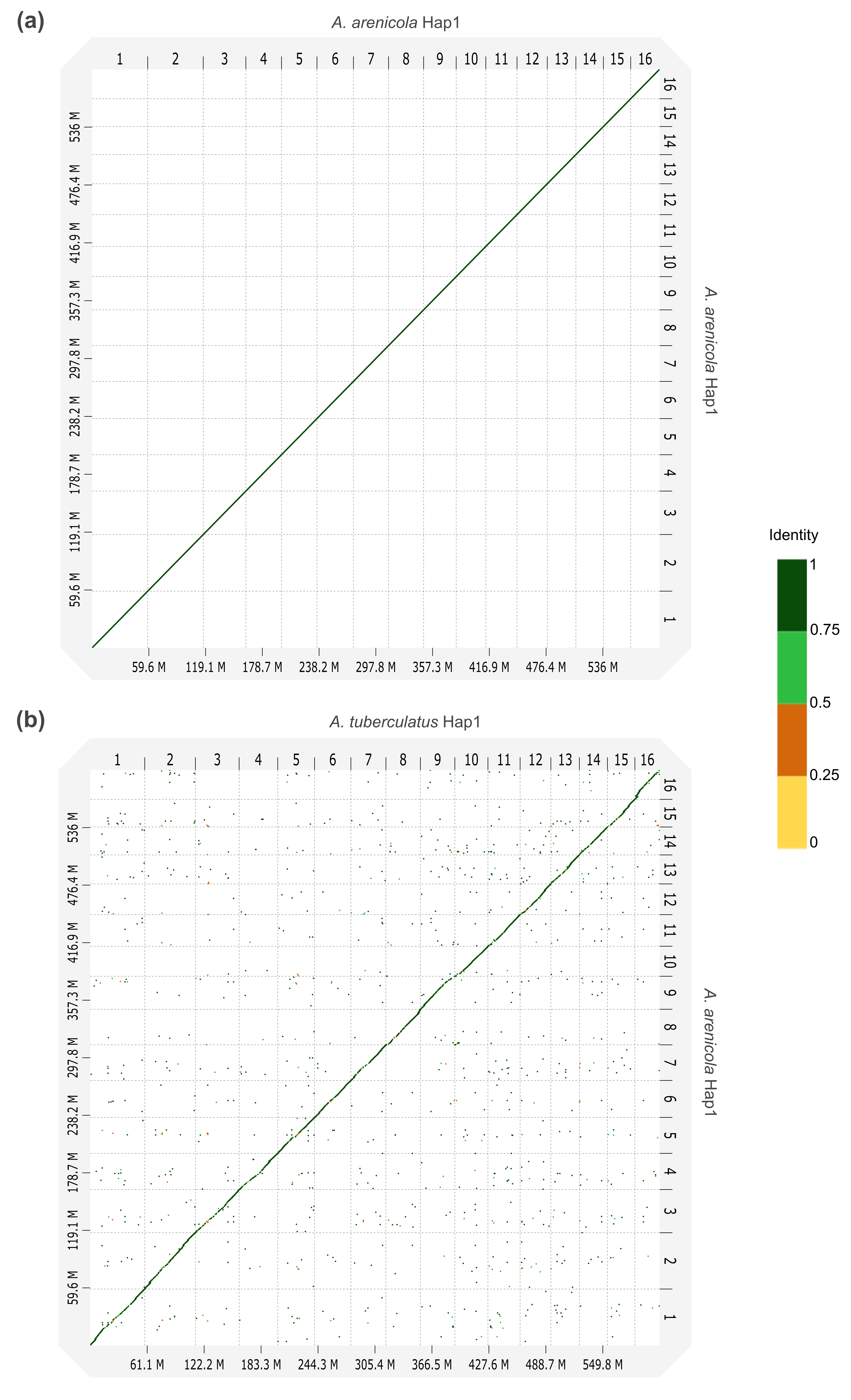

**Figure S5.** Whole-genome pairwise (a) self-alignment of *A. arenicola* Hap1, and (b) alignment between *A. tuberculatus* Hap1 and *A. arenicola* Hap1.

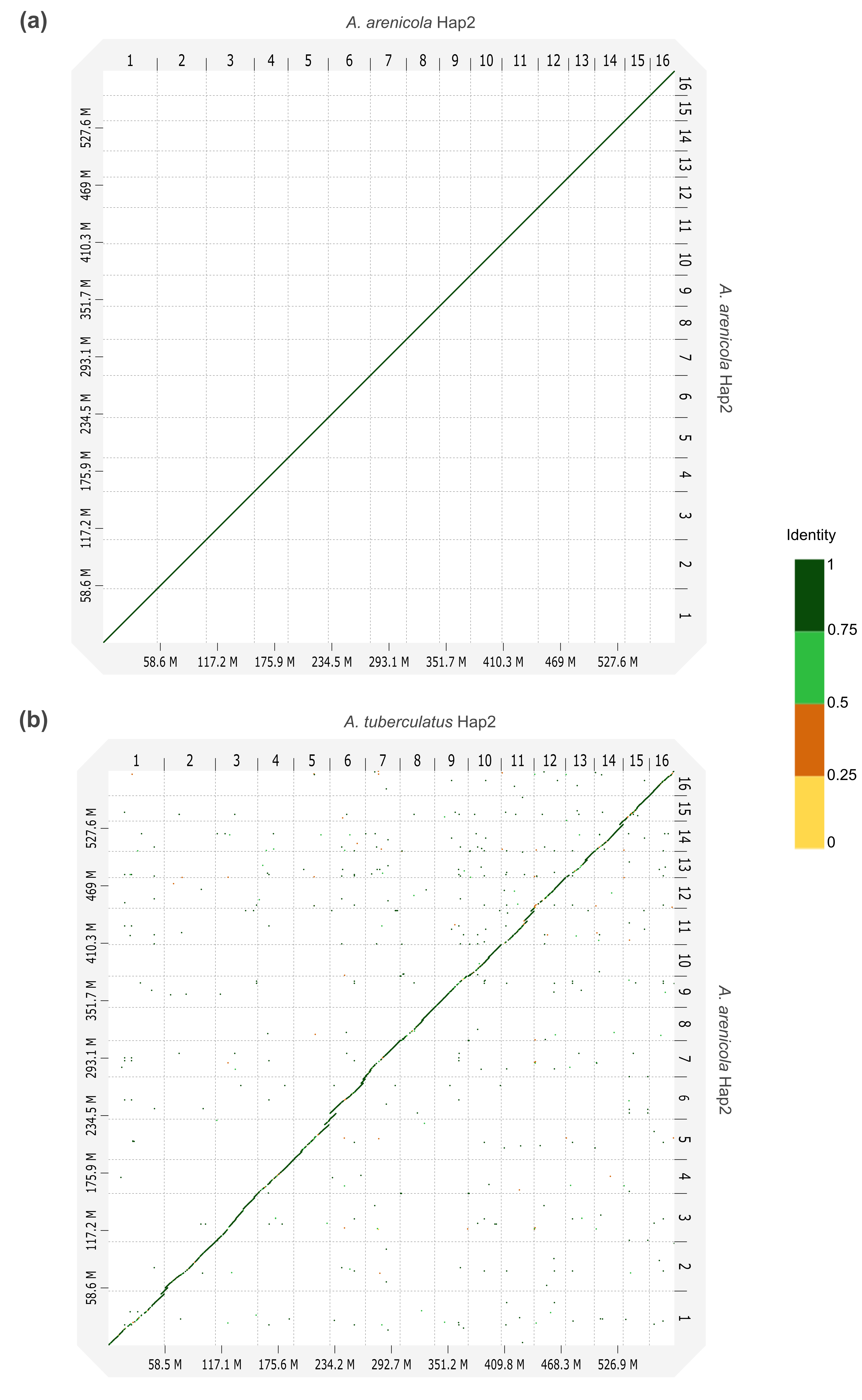

**Figure S6.** Whole-genome pairwise (a) self-alignment of *A. arenicola* Hap2, and (b) alignment between *A. tuberculatus* Hap2 and *A. arenicola* Hap2.

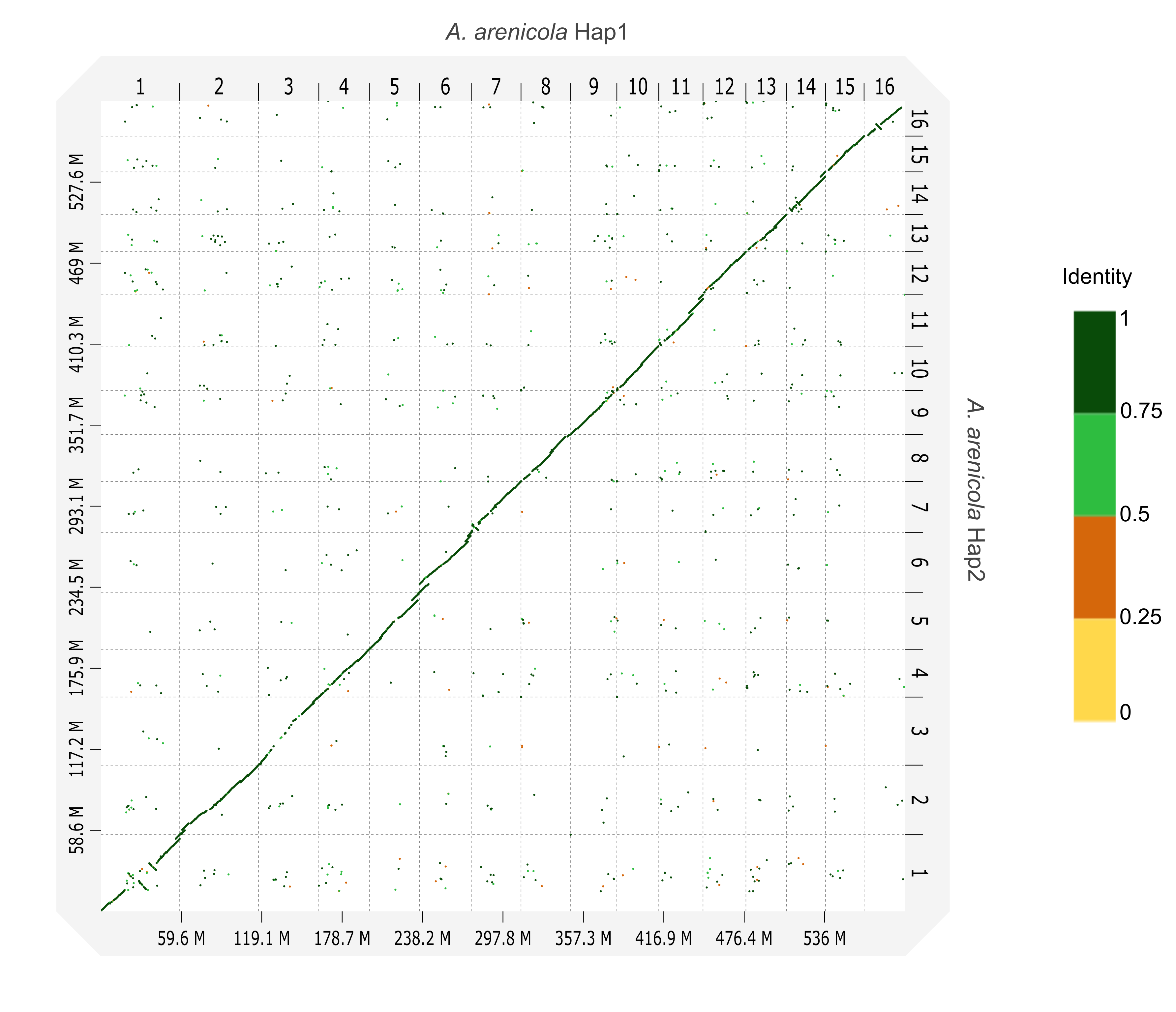

**Figure S7.** Whole-genome pairwise alignment between *A. arenicola* Hap1 and *A. arenicola* Hap2.

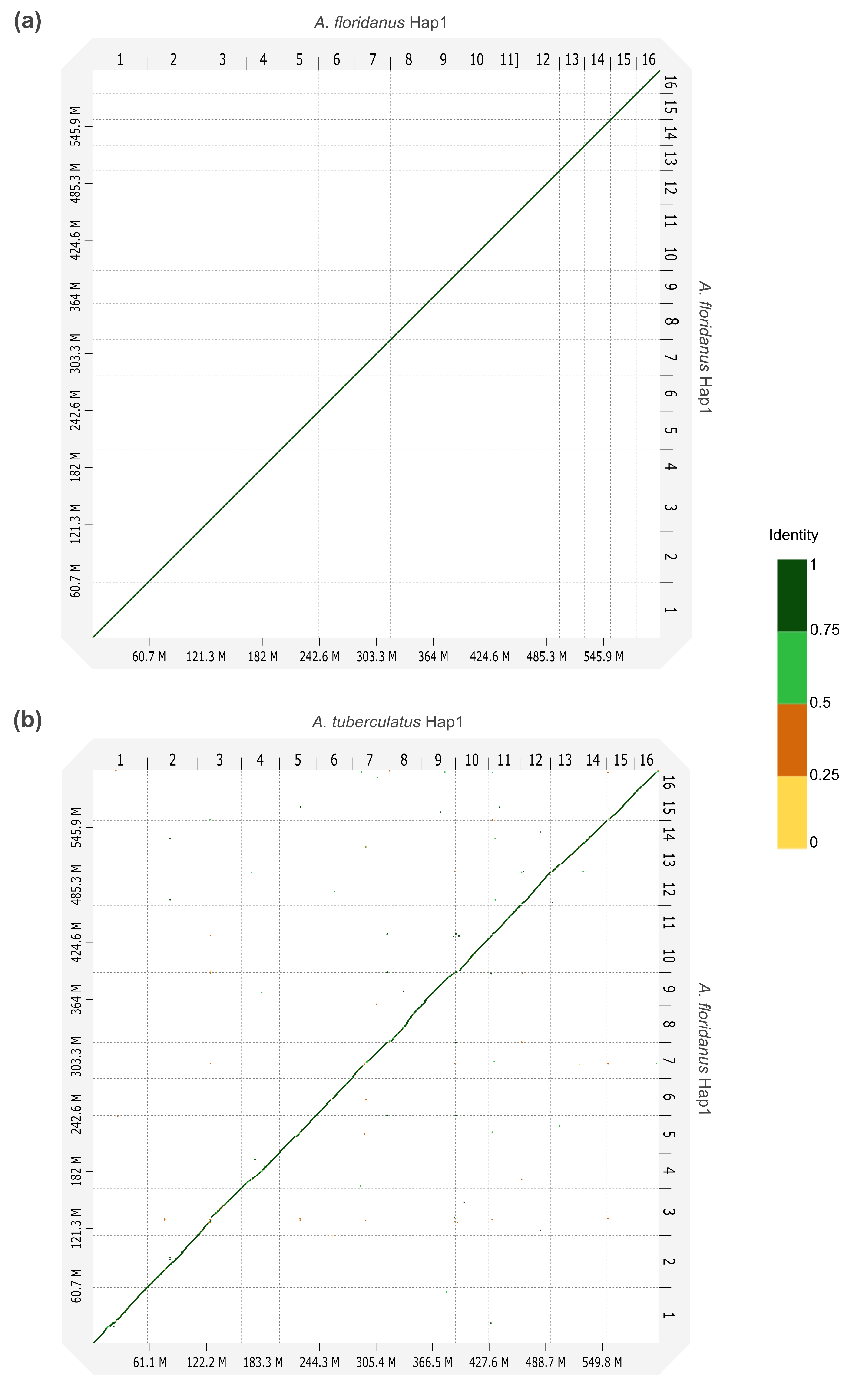

**Figure S8.** Whole-genome pairwise (a) self-alignment of *A. floridanus* Hap1, and (b) alignment between *A. tuberculatus* Hap1 and *A. floridanus* Hap1.

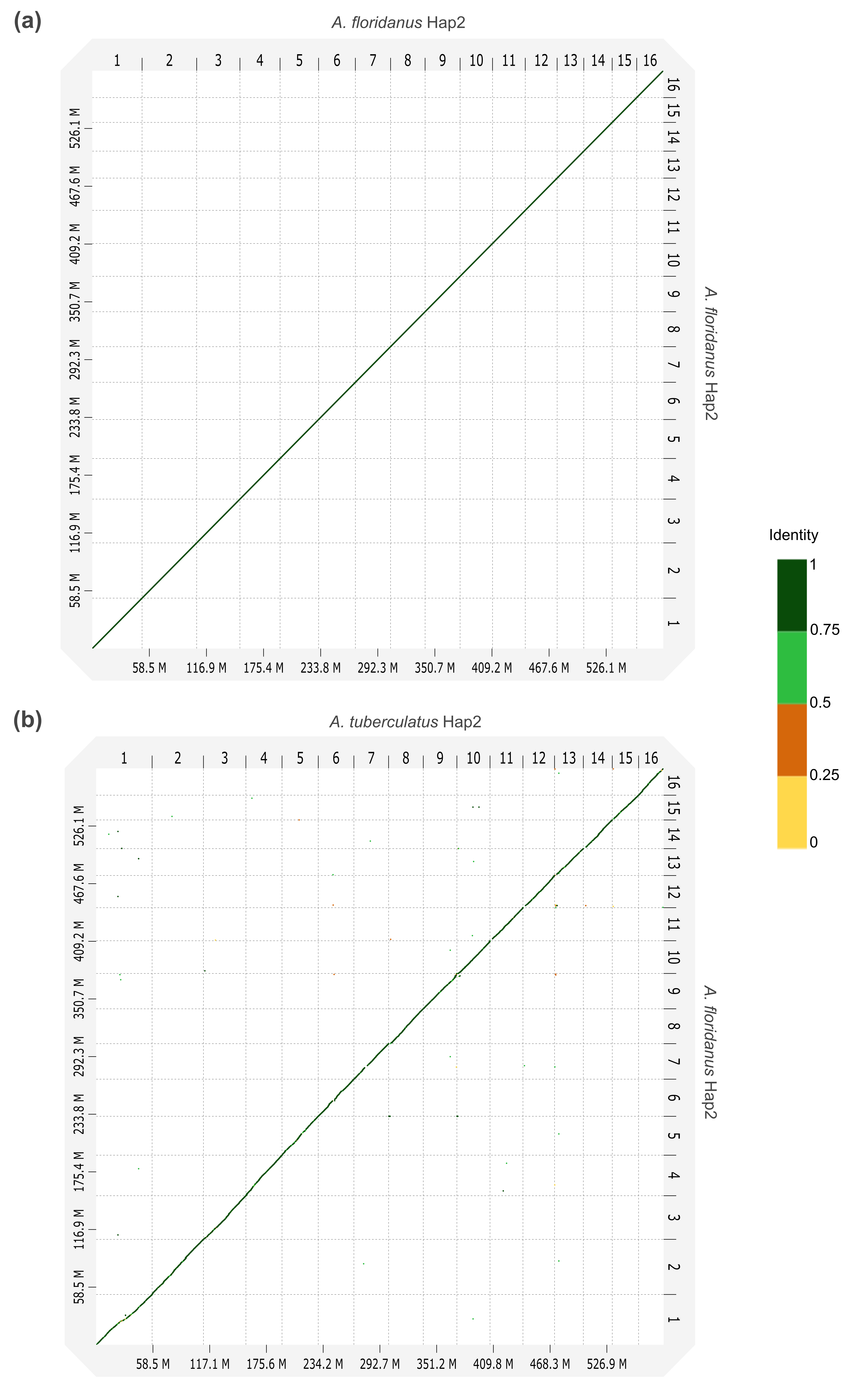

**Figure S9.** Whole-genome pairwise (a) self-alignment of *A. floridanus* Hap2, and (b) alignment between *A. tuberculatus* Hap2 and *A. floridanus* Hap2.

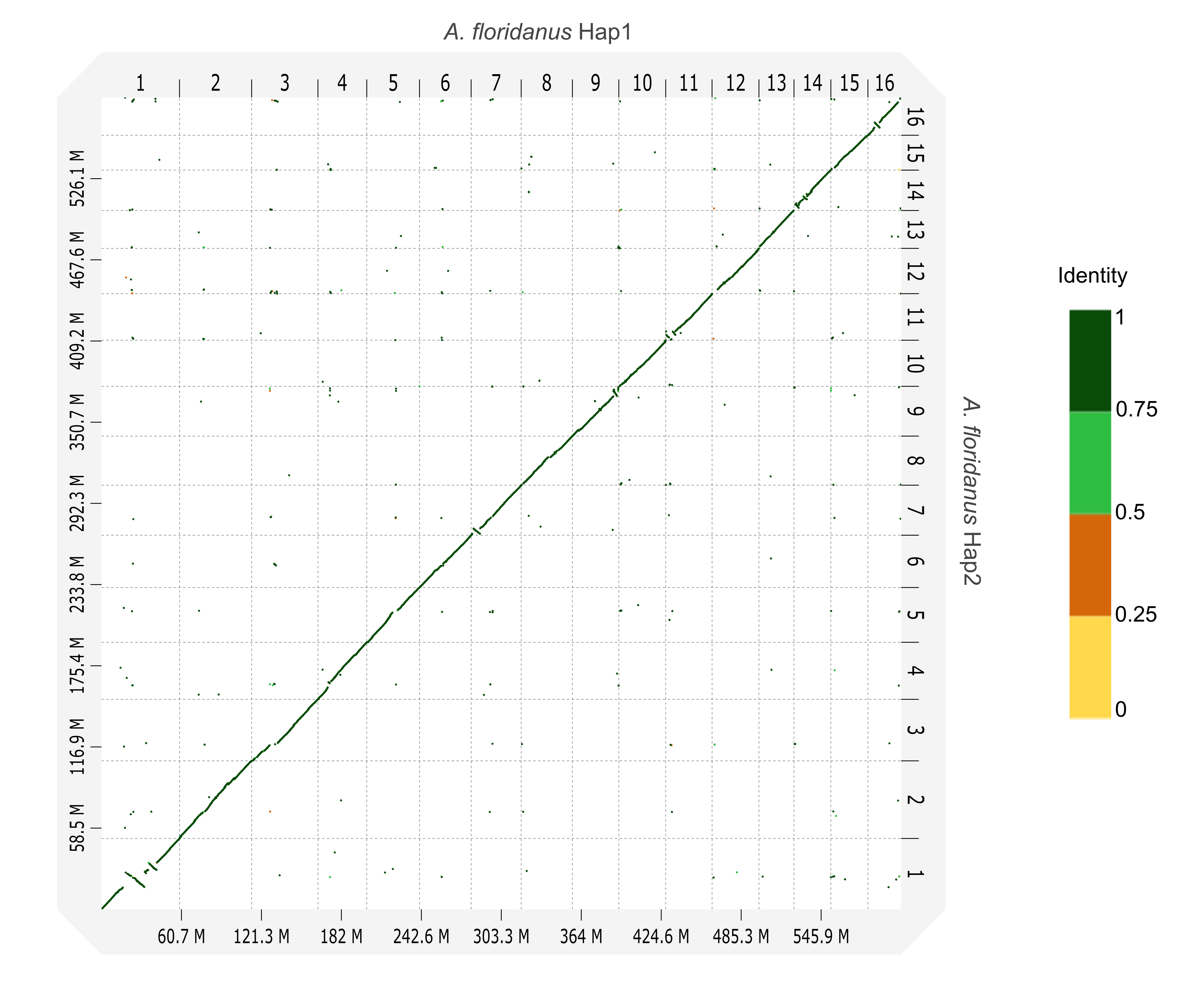

**Figure S10.** Whole-genome pairwise alignment between *A. floridanus* Hap1 and *A. floridanus* Hap2.

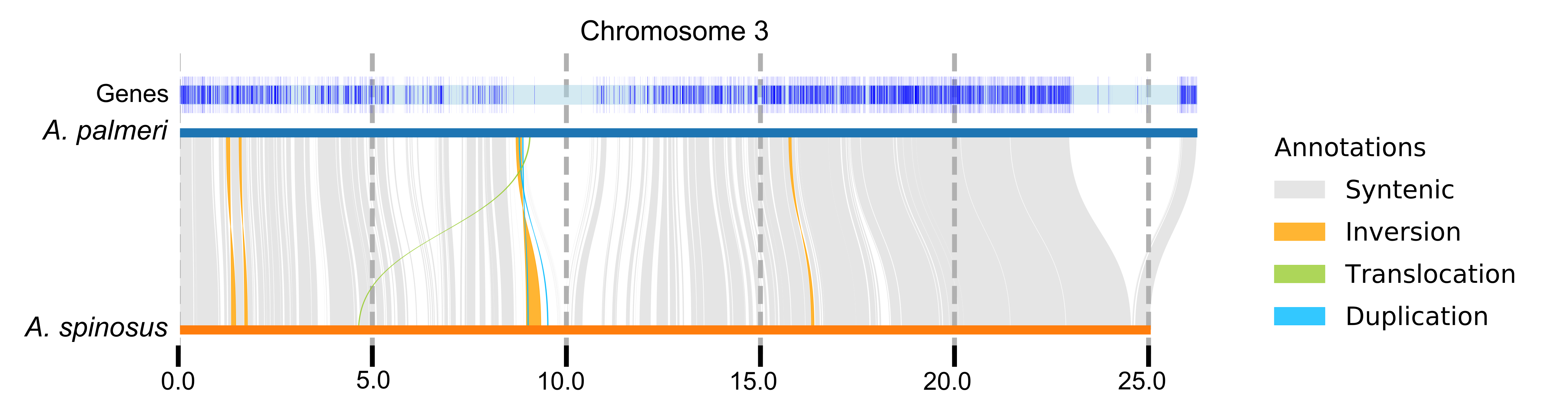

**Figure S11.** Synteny and rearrangement plot (SyRI) between *A. palmeri* Hap1 and *A. spinosus* Hap1, providing a structural variant-level view of the same comparison shown as a whole-genome dotplot in Figure S2a.

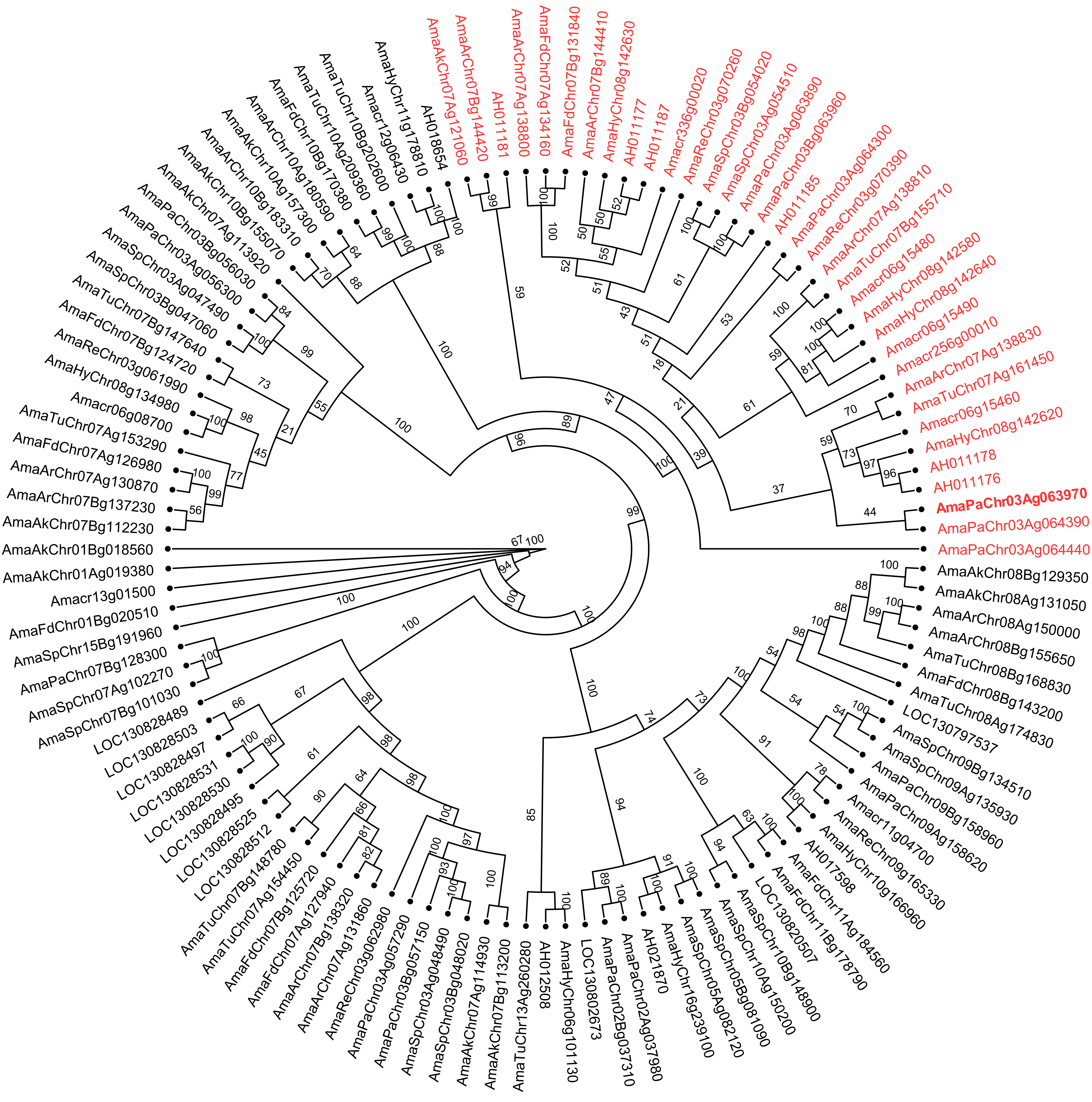

**Figure S12.** Phylogenetic tree of Rf1 protein sequences in *A. palmeri* and its homologs in other *Amaranthus* species (*A. hypochondriacus*, *A. hybridus*, *A. cruentus*, *A. retroflexus*, *A. spinosus*, *A. acanhtochiton*, *A. arenicola*, *A. floridanus*, *A. tuberculatus*, and *A. tricolor*). Taxa indicated in red represent the Rf1 clade while AmaPaChr03Ag063970 in bold is the previously reported Rf1 within the region associated with sex in *A. palmeri*. Values above branches represent ultrafast bootstrap (UFBoot2) support from IQ-TREE3.

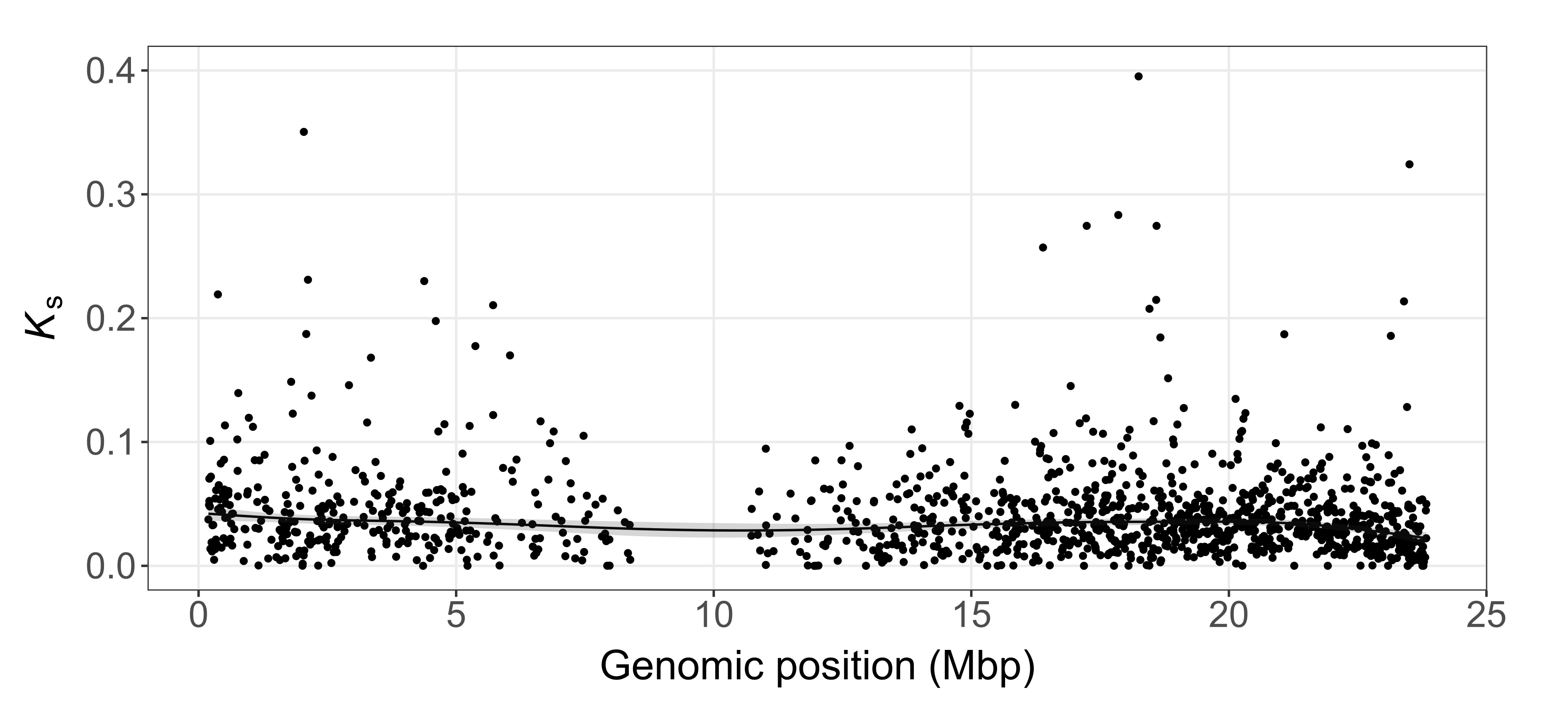

**Figure S13.** Synonymous divergence (Ks) between one-to-one X–Y gametologs along the X chromosome (Chr03) of *A. palmeri*. Robust generalized additive model (GAM) smoothing of Ks was performed on 1,268 gametolog pairs (1:1 orthologs) using penalized regression splines (k = 30 basis functions) and scaled-t error distribution; the fitted smooth (solid line) is shown with its 95% confidence band. The smooth term was significant (p ≤ 0.0003) but explained only 1.3% of Ks variance (adjusted R^2^), indicating weak spatial structure with no discrete divergence peak consistent with a classical multi-stratum architecture.

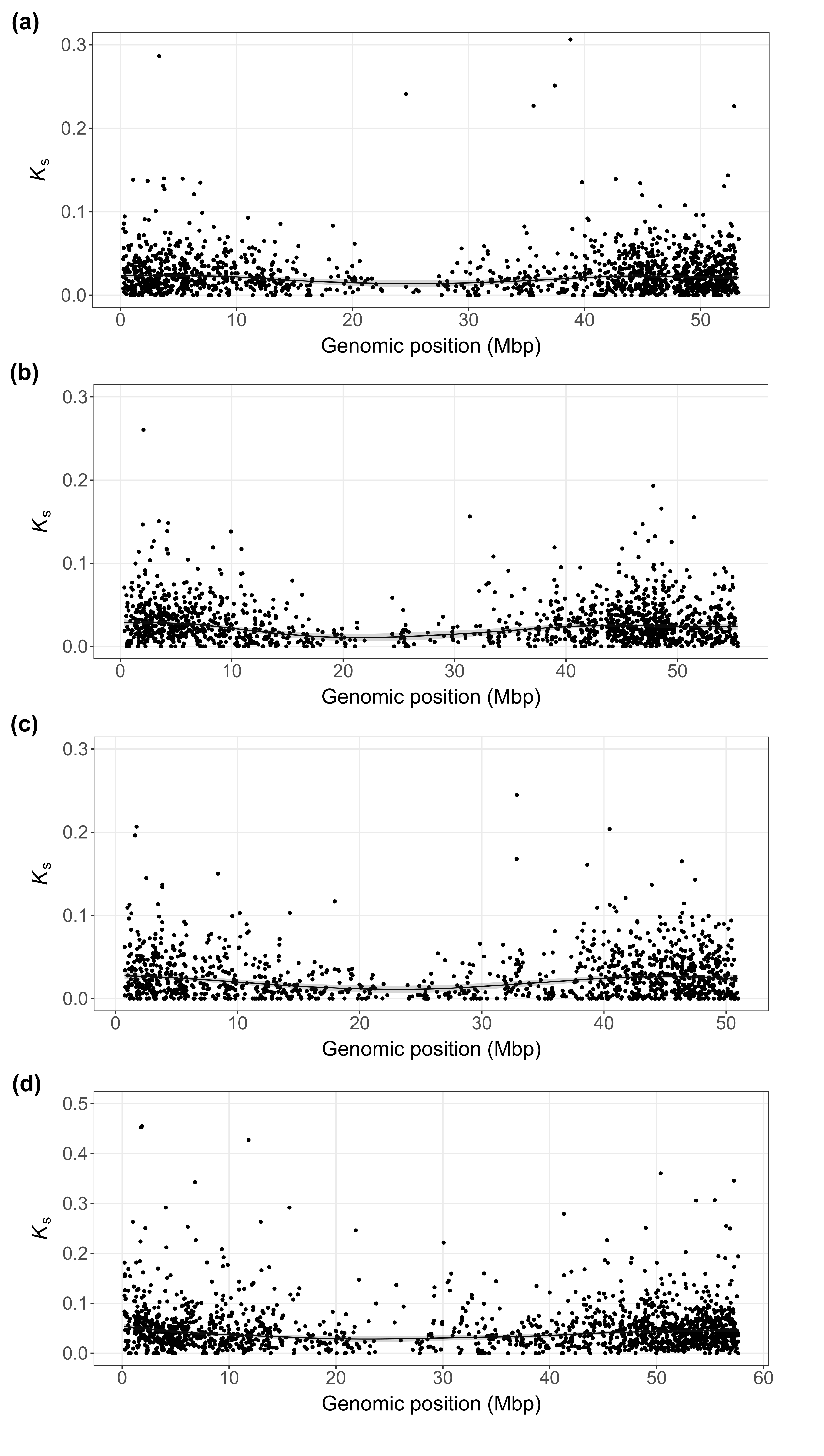

**Figure S14.** Synonymous divergence (Ks) between one-to-one X–Y gametologs along the X chromosome (Chr01) of *A. acanthochiton* (a), *A. arenicola* (b), *A. floridanus* (c), and *A. tuberculatus* (d). Robust generalized additive model (GAM) smoothing of Ks was performed using penalized regression splines (k = 30 basis functions) and a robust scaled-t error distribution; each fitted smooth (solid line) is shown with its 95% confidence band. Datasets comprised 1,575 gametolog pairs in *A. acanthochiton* (a), 1,483 in *A. arenicola* (b), 1,271 in *A. floridanus* (c), and 1,710 in *A. tuberculatus* (d). Smooth terms were significant in all species (p ≤ 0.0002) but explained only a small fraction of Ks variance [adjusted R^2^: 0.5% (a), 2.8% (b), 3.2% (c), 1.1% (d)], indicating weak spatial structure with no discrete divergence peaks consistent with a classical multi-stratum architecture.

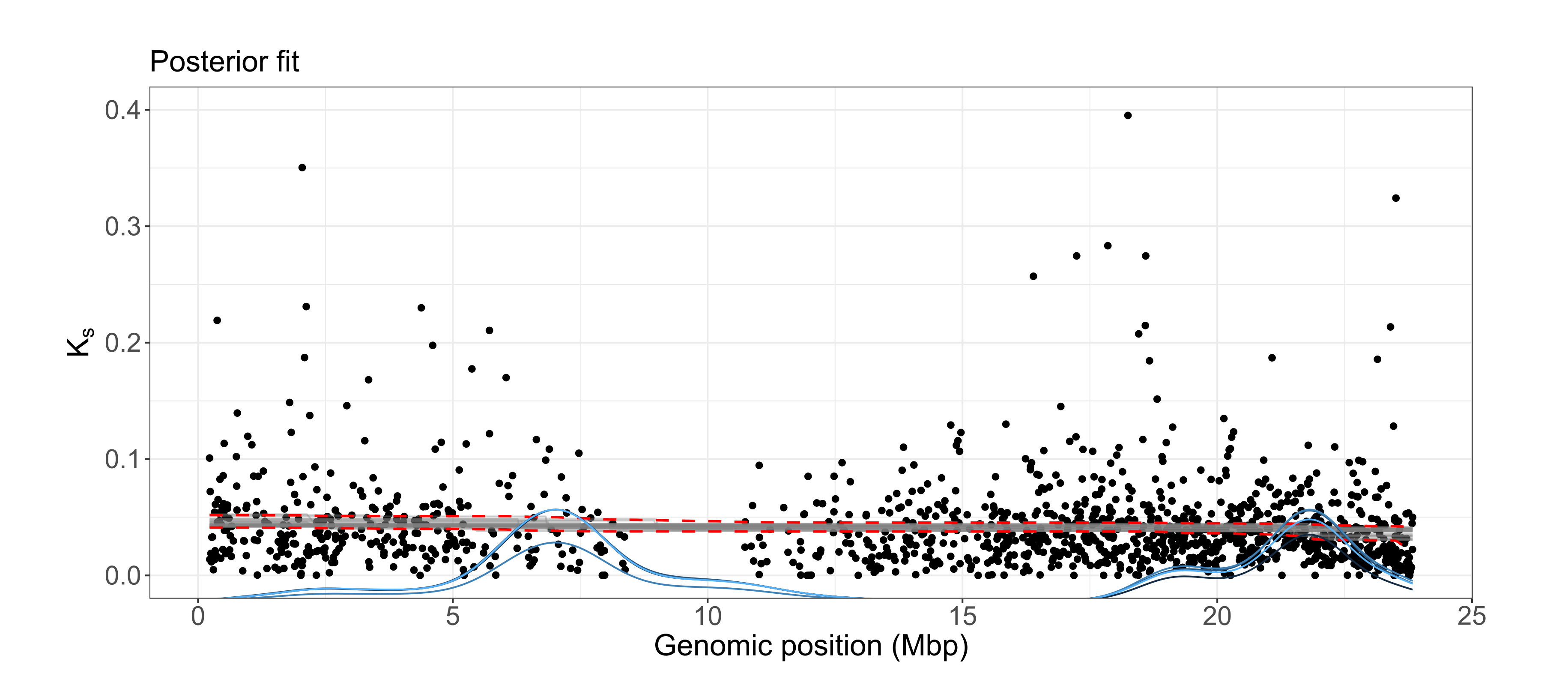

**Figure S15.** Distribution of synonymous divergence (Ks) values from one-to-one orthologs along X chromosome (Chr03) of *A. palmeri*. Changepoint analysis of Ks along the X chromosome (Chr03) using a set of 1,268 gametologs with 1:1 orthologs. The gray lines represent the posterior fitted draws. Red band represents the posterior quantile summary of the fitted curve. The blue line represents the posterior distribution of changepoint location(s), scaled into the plot’s y-range.

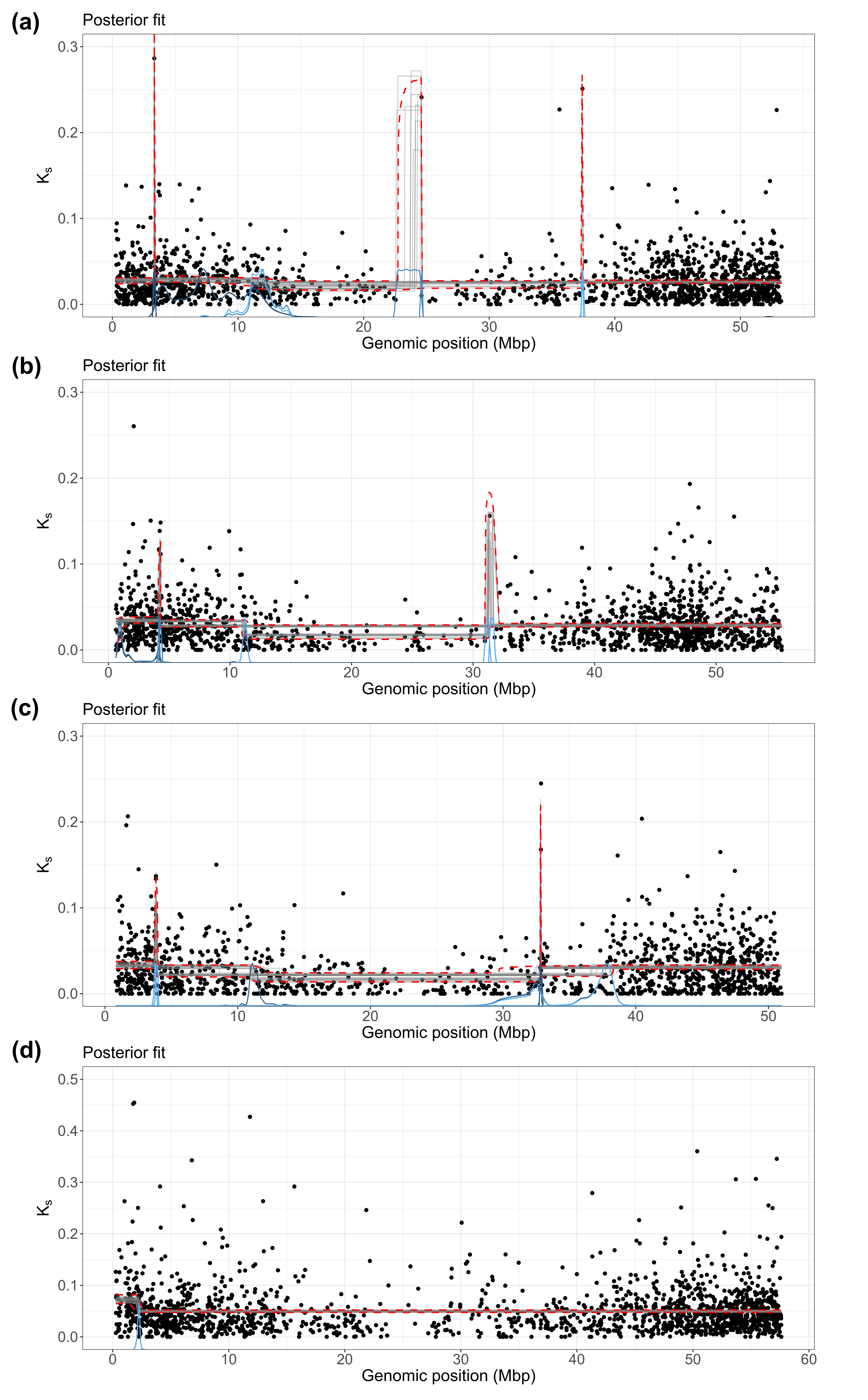

**Figure S16.** Distribution of synonymous divergence (Ks) values from one-to-one orthologs along the X chromosome (Chr01) of *A. acanthochiton* (a), *A. arenicola* (b), *A. floridanus* (c), and *A. tuberculatus* (d). Changepoint analysis of Ks along the X chromosome (Chr01) using a set of 1,575 gametologs with 1:1 orthologs in *A. acanthochiton* (a), 1,476 gametologs in *A. arenicola* (b), 1,267 gametologs in *A. floridanus* (c), and 1,710 gametologs in *A. tuberculatus* (d). The gray lines represent the posterior fitted draws. Red band represents the posterior quantile summary of the fitted curve. The blue line represents the posterior distribution of changepoint location(s), scaled into the plot’s y-range. Panels (b) and (c) are shown for completeness but excluded from the interpretation presented in the main text.

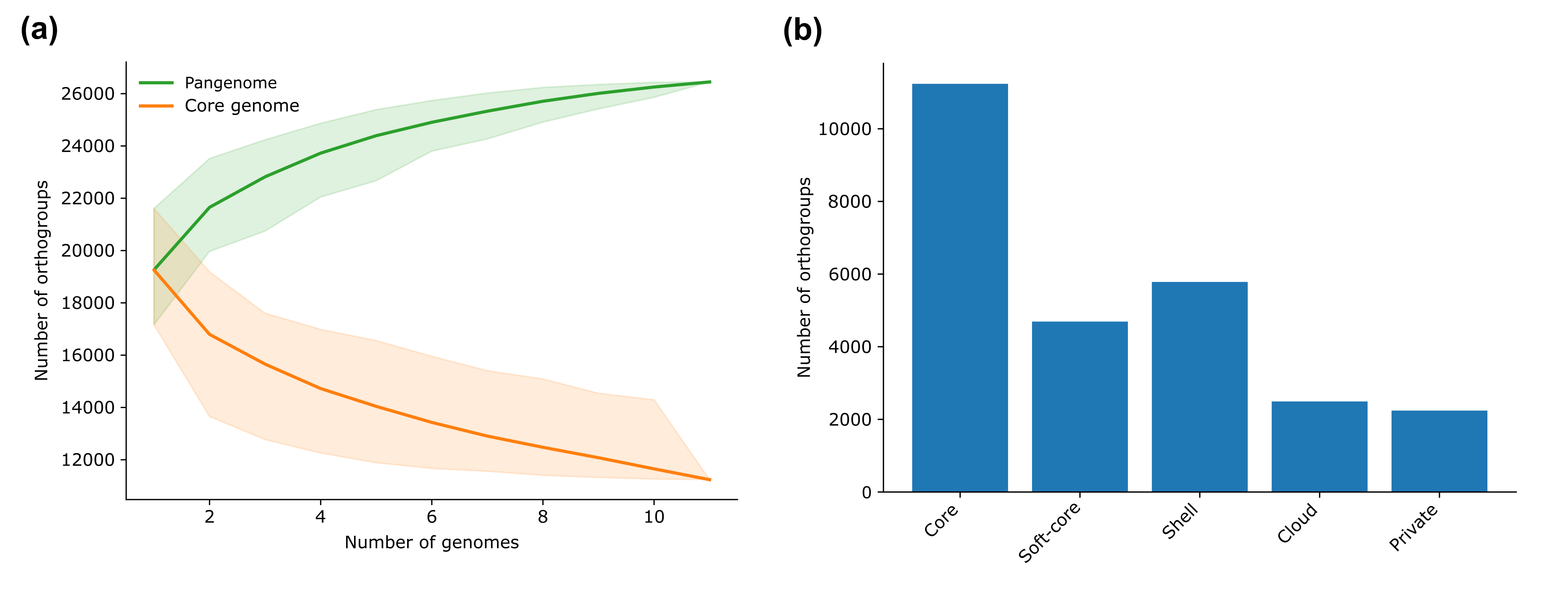

**Figure S17.** Pangenome analysis of 11 *Amaranthus* species genomes. (a) Accumulation curves of orthogroups (gene families) of the pangenome (green) and core-genome (orange). Shaded regions show the 2.5th – 97.5th percentile range across random permutations of genome order to determine how quickly the pan/core stabilizes (i.e. heterogeneity) as genomes are added. (b) bar plot of the pangenome compartments.
